# Spatially resolved whole transcriptome profiling in human and mouse tissue using Digital Spatial Profiling

**DOI:** 10.1101/2021.09.29.462442

**Authors:** Stephanie M. Zimmerman, Robin Fropf, Bridget R. Kulasekara, Maddy Griswold, Oliver Appelbe, Arya Bahrami, Rich Boykin, Derek L. Buhr, Kit Fuhrman, Margaret L. Hoang, Quoc Huynh, Lesley Isgur, Andrew Klock, Alecksandr Kutchma, Alexa E. Lasley, Yan Liang, Jill McKay-Fleisch, Jeffrey S. Nelson, Karen Nguyen, Erin Piazza, Aric Rininger, Daniel R. Zollinger, Michael Rhodes, Joseph M. Beechem

## Abstract

Emerging spatial profiling technology has enabled high-plex molecular profiling in biological tissues, preserving the spatial and morphological context of gene expression. Here we describe expanding the chemistry for the Digital Spatial Profiling platform to quantify whole transcriptomes in human and mouse tissues using a wide range of spatial profiling strategies and sample types. We designed multiplexed *in situ* hybridization probe pools targeting the protein-coding genes in the human and mouse transcriptomes, hereafter referred to as the human or mouse Whole Transcriptome Atlas (WTA). We validated the human and mouse WTA using cell lines to demonstrate concordance with orthogonal gene expression profiling methods in profiled region sizes ranging from ~10-500 cells. By benchmarking against bulk RNAseq and fluorescence in situ hybridization, we demonstrate robust transcript detection possible down to ~100 transcripts per region. To assess the performance of WTA across tissue and sample types, we applied WTA to biological questions in cancer, molecular pathology, and developmental biology. We show that spatial profiling with WTA can detect expected spatial gene expression differences between tumor and tumor microenvironment, identify spatial disease-specific heterogeneity in gene expression in histological structures of the human kidney, and comprehensively map transcriptional programs in anatomical substructures of nine organs in the developing mouse embryo. Digital Spatial Profiling technology with the WTA assays provides a flexible method for spatial whole transcriptome profiling applicable to diverse tissue types and biological contexts.

## Introduction

The organization of tissues and organs is complex and spatial relationships between cells and structures are key to their development, normal functioning, and pathophysiology. Recently, several methods have emerged for multiplexed spatial profiling of RNA or proteins, leading to discoveries in oncology, infectious disease, developmental biology, and other fields (Rao et al. 2021; Brady et al. 2021; Butler et al. 2021; Pelka et al. 2021; Jerby-Arnon et al. 2021; Desai et al. 2020; Rendeiro et al. 2021; Merritt et al. 2020). Existing spatial gene expression platforms operate at a range of plex and with differing profiling strategies. Sequencing-based methods capture transcripts in an unbiased manner and are capable of whole transcriptome coverage. For example, laser capture microdissection is a method for physically separating cells and structures of interest within a tissue, which can then be subjected to a variety of molecular profiling methods including RNAseq (Emmert-Buck et al. 1996; Espina et al. 2006). Other sequencing-based methods such as Slide-Seq and Spatial Transcriptomics capture polyadenylated mRNAs across pre-patterned barcoded spot arrays (Vickovic et al. 2019; Ståhl et al. 2016; Stickels et al. 2021). An advantage of these methods is that they can provide unbiased coverage of the transcriptome. However, one disadvantage is that RNAseq via poly-A capture can be dominated by highly expressed genes.

Imaging-based methods, such as multiplexed error-robust fluorescence in situ hybridization (MERFISH), fluorescent in situ sequencing (FISSEQ), and Sequential barcoded Fluorescence i*n situ* Hybridization (seqFISH) (Chen et al. 2015; Lee et al. 2015; Xia et al. 2019), use rounds of sequential hybridization and imaging to resolve transcripts at single cell or subcellular resolution. Some imaging methods have demonstrated detection of up to 10,000 targets, but most experiments have been limited to lower plex in the hundreds of targets (Eng et al. 2019; Xia et al. 2019).

Digital Spatial Profiling (DSP) is a recently developed platform for multiplexed spatial RNA or protein expression profiling in user-defined regions of interest (Merritt et al. 2020). DSP relies on affinity reagents (probes for RNA and antibodies for protein detection) attached to indexing oligonucleotide tags with a UV-photocleavable linker. The probes or antibodies are hybridized to a slide-mounted tissue sample that is also stained with fluorescent antibodies or probes to aid in the identification of features of interest. The tissue is imaged using fluorescence microscopy and UV light is projected onto the region to be profiled, called areas of illumination (AOIs), to release the oligo tags from that region. The liberated tags are collected and counted using the nCounter® system or by next-generation sequencing (NGS). In the first demonstration of the DSP technology, 44 proteins and 84 genes were multiplexed using nCounter, and 1,412 genes were profiled by NGS readout (Merritt et al. 2020).

Here we report the expansion of the DSP RNA profiling technology to measure the expression of >99.5% and >98.2% of protein-coding genes of the human or mouse transcriptome, respectively, with a small number of very highly expressed genes intentionally removed to provide better coverage of low expressing transcripts. For all annotated genes, we designed in situ hybridization (ISH) probes with a barcoded UV-cleavable tag that can be read out by NGS. The probes for each species were pooled to create the human and mouse Whole Transcriptome Atlases (WTAs). In this study, we aim to characterize the technical performance of the human and mouse WTA across a range of region sizes and profiling strategies, and demonstrate applications in diverse tissue contexts.

## Results

### Design of multiplexed probes targeting the human and mouse whole transcriptomes

The human or mouse WTA consists of species-specific ISH probes designed to target the protein-coding genes of the human or mouse transcriptome. The probes contain three functional regions: an RNA-targeting region, a UV-photocleavable linker, and an indexing sequence designed to be read out by NGS. The indexing sequence contains a Unique Molecular Identifier (UMI), a barcode sequence that identifies the probe, and primer binding sequences for amplification and subsequent readout by standard NGS workflows (Supplemental Fig. S1). The probe identification barcodes were designed to have a minimum Hamming distance of ≥2 between barcodes.

We designed 18,815 human and 20,175 mouse probes targeting >99.5% of annotated protein-coding genes in human and >98.2% of annotated protein coding genes in mouse (Supplemental Table S1). To reduce sequencing requirements and optimize readout efficiency by NGS, probes targeting mitochondrially encoded genes and an additional 10 human and 2 mouse highly expressed nuclear-encoded genes were intentionally removed (see Methods). Mouse WTA also includes probes targeting 17 commonly used transgenes. We additionally designed 139 negative control probes in human WTA and 210 negative control probes in mouse WTA against synthetic sequences from the External RNA Controls Consortium (ERCC) set (Baker et al. 2005). The ERCC sequences have the same properties as mammalian sequences but without similarity to any known transcripts. This was confirmed by BLAST comparison to each transcriptome for all selected negative sequences.

RNA-targeting regions range in size from 35-50 nucleotides and were selected based on an iterative design process that considers thermodynamic profile, splice isoform coverage, potential for cross-hybridization with other transcripts, and potential for intramolecular interactions between probes within an assay (see Methods). Probes were synthesized individually and pooled, and the pools were sequenced to ensure that 100% of designed probes were present and that the coefficient of variation of probe concentration was less than 20%.

As WTA contains a single probe per gene, we assessed the consequences of this design choice by comparing human WTA to a smaller probe pool targeting 1,812 human genes with 5 probes per target. We compared counts in matched 200 μm AOIs in formalin-fixed paraffin embedded (FFPE) tonsil tissue, and found that counts from the single WTA probe were well correlated to the mean count of the five probes for the same target (median R = 0.83), as well as a randomly selected single probe (median R = 0.73) (Supplemental Fig. S2). These results validate that a single probe is generally sufficient to accurately quantify gene expression.

### WTA data are reproducible and well correlated with RNAseq and RNA FISH in cell lines

We first benchmarked the performance of the human and mouse WTAs in homogeneous FFPE cell pellet arrays (CPAs) to test reproducibility and compare to orthogonal methods of measuring gene expression. Because DSP allows flexible selection of the areas of illumination on the tissue (AOIs), it is possible to profile regions of various sizes ranging from <10 cells to thousands of cells. As there is a tradeoff between the number of cells profiled and signal, we benchmarked the performance of WTA in AOIs ranging from 50-400 μm diameter circles in human and mouse FFPE CPAs containing 11 cell lines each (Fig. 1A). In cell pellets, 50 μm diameter AOIs contained an average of 12 cells in human cell lines and 13 cells in mouse cell lines, while 400 μm diameter AOIs contained an average of 480 and 505 cells in human and mouse, respectively. Counts were highly reproducible between two independent experiments for all cell lines and AOI sizes tested (R = ~0.75 for 50 μm AOIs, and ~0.95 for 400 μm AOIs for all genes) for both human and mouse WTAs (Fig. 1B).

**Figure 1.**
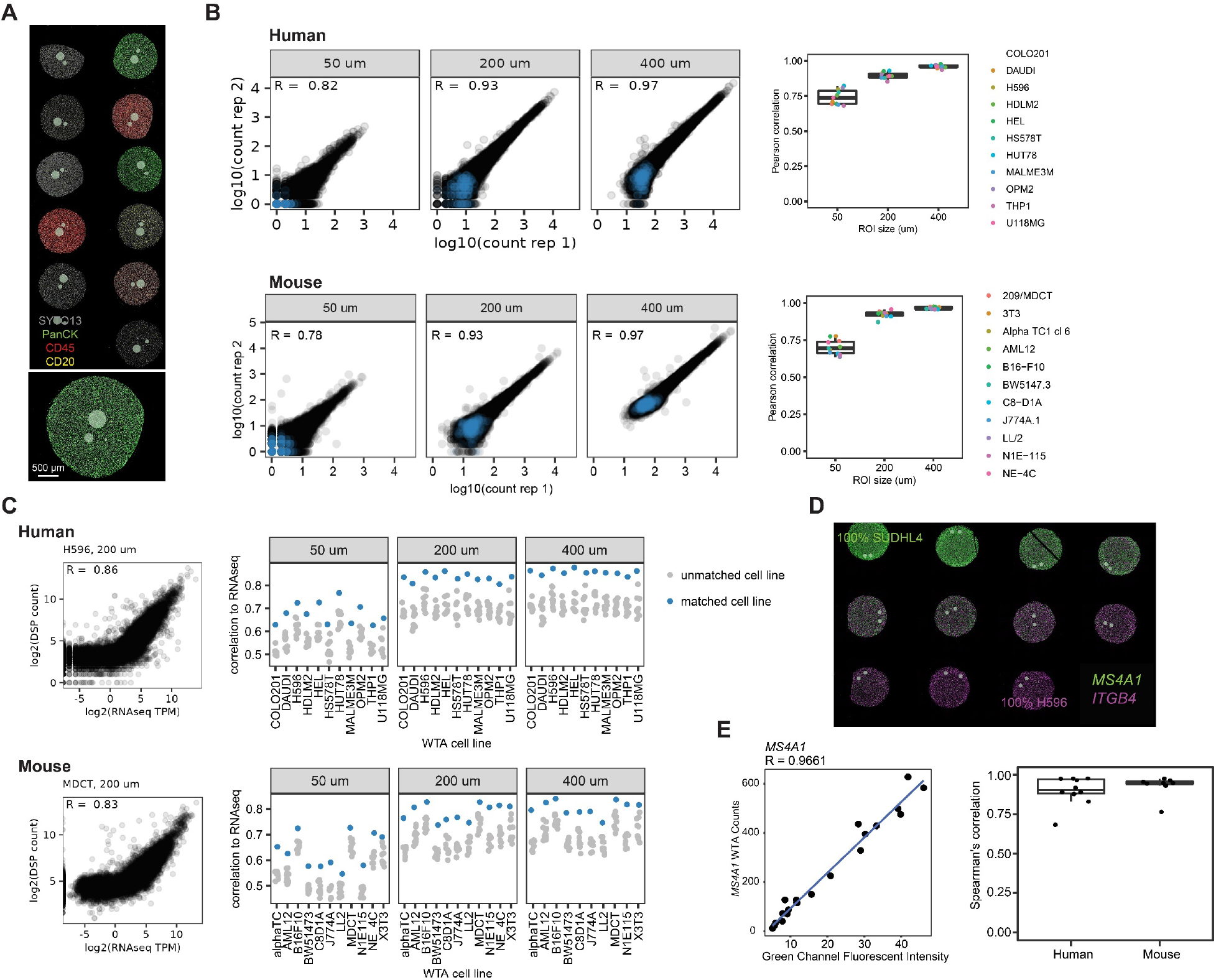
Human and mouse WTA data are reproducible and correlated with RNAseq and RNA FISH. **A.** Representative image of the AOI size titration experiment. Circular AOIs 50 μm, 200 μm, and 400 μm in diameter were placed on each cell line of an 11-core human or mouse FFPE cell pellet array (human shown, stained with antibodies against CD3, CD45 and pan-cytokeratin (PanCK), and SYTO13 nuclear stain). **B.** Reproducibility of WTA counts from two replicate experiments. Left: scatterplots of log2-transformed raw counts from one representative human or mouse cell line (HUT78 for human, 3T3 for mouse) at each AOI size from each replicate. Negative control probes are shown in blue and target probes in black. Right: Pearson correlation coefficients of log2-transformed raw counts between replicates for each cell line and AOI size. **C.** Left: scatterplots of WTA counts vs RNAseq transcripts per million (TPM) from the same cell line for one representative human or mouse cell line in a 200 μm AOI. Right: Spearman’s correlation of WTA counts compared to RNAseq of each cell line profiled in this experiment. For each AOI, the matching cell line is shown in blue and all other cell lines in grey. **D.** Representative image of the cell line titration experiment. Cell pellets contained one cell line titrated into the other at a variable ratio. Cells were stained with RNAscope probes against two genes specifically expressed in each of the two cell lines (ITGB4 expressed in H596 cells and MS4A1 expressed in SUDHL4 cells in the image shown). Grey circles show profiled AOIs. **E.** Left: Representative scatterplot comparing WTA counts for MS4A1 to RNAscope fluorescence intensity for the same gene across cell pellets. Right: Spearman’s correlation of WTA counts compared to RNA FISH fluorescence intensity for each gene profiled in this experiment.

We asked whether we could correctly classify these cell lines based on WTA signal, comparing to bulk RNAseq of the same set of cell lines. Bulk RNAseq data were either generated for this study or acquired from publicly available data from the Cancer Cell Line Encyclopedia (CCLE) project (Ghandi et al. 2019) (see Methods). Using all genes in the WTA panels, we found that classification was 100% accurate at all AOI sizes. For all 11 human and 11 mouse cell lines, the correct matching cell line had the highest correlation coefficient between WTA and bulk RNAseq. Correlation coefficients with the matching cell line were ~0.7 in 50 μm diameter circle AOIs and increased to >0.8 in 400 μm diameter circle AOIs, and were similar for human and mouse WTA (Fig. 1C). To test the effect of gene expression level on cell line classification, we compared WTA counts to RNAseq subset to the lowest and highest quartile of expressed genes, defined as genes >1 transcript per million (TPM) in RNAseq. Discrimination between the correct matching and non-matching cell lines was maintained for the lowest quartile of expressed genes in 200 μm and 400 μm AOIs, but was reduced for 50 μm AOIs (Supplemental Fig. S2), suggesting that very lowly expressed genes are not as well quantified in very small AOIs.

We next tested whether we could accurately quantitate gene expression with WTA. For these experiments, we used a mixed-proportion FFPE (human) or fixed frozen (mouse) CPA with one cell line titrated into another in 10% increments. We selected 9 human genes and 8 mouse genes that are highly expressed in one cell line (>100 TPM in bulk RNAseq) and not expressed in the other (<1 TPM in bulk RNAseq) (Supplemental Fig. S3). In the CPA, this creates a gradient of gene expression levels across the different cell pellets that spans the range of very lowly to very highly expressed genes. For each gene, WTA signal was compared to FISH signal using RNAscope probes (Wang et al. 2012) (Fig. 1D). We found that WTA and FISH signals were highly correlated for all genes tested, with an average Pearson correlation coefficient of 0.90 for human and 0.93 for mouse (Fig. 1E, Supplemental Fig. S3). These results indicate that WTA can accurately quantify gene expression across the biological range of gene expression.

### Sensitivity, specificity, and limit of detection of WTA in different sized AOIs

We next investigated WTA’s sensitivity and specificity to detect gene expression above background in different sized AOIs. We used the distribution of signal from the negative control probes to estimate the background level of non-specific binding and set a limit of detection (LOD) specific to each AOI. Counts from individual negative probes are moderately correlated between replicate experiments, suggesting that the variance in negative probe signal is due to both sequence-specific and non-specific effects (Supplemental Fig. S4). Both target and negative probe counts increase with AOI area and scale with each other across different cell lines (Supplemental Fig. S4), highlighting the importance of empirically measuring the non-specific background for each AOI.

To determine a cutoff for calling a gene expressed, we calculated sensitivity and specificity at different thresholds above background using genes with RNAseq TPM > 1 as the true set of expressed genes. With an increasing threshold, specificity increases but sensitivity decreases; discrimination improves with increasing AOI size and is similar between human and mouse WTA (Fig. 2A). We found that selecting an LOD threshold of 2 standard deviations above the geometric mean of negative probes reliably achieves a specificity of >95% at all ROI sizes and in both panels. At this LOD threshold, sensitivity was 50% in 50 μm diameter-circle AOIs, 68% in 200 μm AOIs, and 81% in 400 μm AOIs for human WTA, and 48% in 50 μm AOIs, 66% in 200 μm AOIs, and 75% in 400 μm AOIs for mouse WTA. Overall, we detect an average of ~6000 genes above background per AOI in 50 μm AOIs, and ~9000 genes in 400 μm AOIs (Fig. 2B).

**Figure 2.**
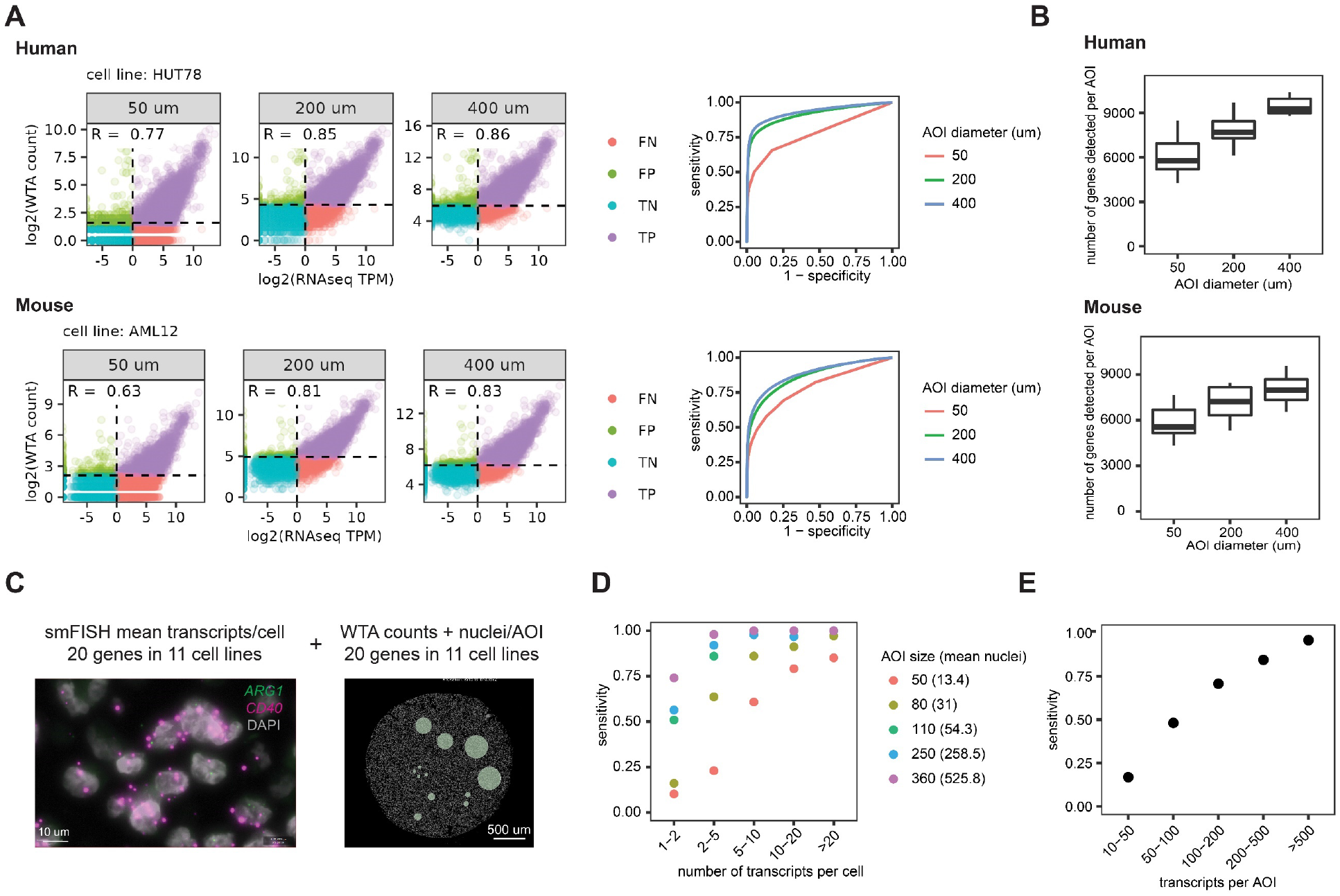
WTA has high sensitivity and can detect genes at a range of expression levels depending on AOI size. **A.** Left: Scatterplots comparing WTA counts to RNAseq for one representative cell line at each AOI size, colored by whether the gene is detected above the expression threshold in WTA and in RNAseq. Dashed lines indicate thresholds for calling a gene “expressed” as 2 standard deviations above the geometric mean of negative probes for WTA, and TPM >1 for RNAseq. TP = true positive, FP = false positive, TN = true negative, and FN = false negative. Right: Receiver-operator curves demonstrating the sensitivity and specificity of WTA at different expression thresholds using genes with RNAseq TPM >1 as the true set of expressed genes. **B.** Number of genes per AOI above the expression threshold of 2 SD above the mean negative probe count at each AOI size. **C.** Representative images of the experiment to determine the sensitivity of human WTA relative to absolute transcript number. Left: RNAscope image of two genes in one cell line of the 20 genes in 11 cell lines quantified in this experiment. Right: DSP image of one cell line with an AOI size titration. **D.** Sensitivity of WTA at different AOI sizes for genes in different gene expression bins as measured by RNAscope. Genes ≥ 1 transcript per cell were considered expressed. **E.** Sensitivity of WTA for genes binned by transcripts per AOI, calculated using transcripts per cell quantified by RNAscope and the number of cells in each AOI.

To determine the LOD of WTA relative to absolute transcript number, we integrated human WTA FFPE CPA data with RNAscope experiments in which we counted the absolute number of transcripts per cell for 20 genes in 11 cell lines (Fig. 2C and Supplemental Fig. S5) (Wang et al. 2012). These genes spanned a range of expression levels across different cell lines, from a mean count of 0 to 45 transcripts per cell (corresponding to 0-1200 TPM in RNAseq) (Supplemental Fig S5). Gene expression levels as measured by RNAscope were well correlated between replicate experiments and well correlated with RNAseq for genes above 1 TPM. Below 1 TPM, all genes had a mean expression of less than 1 transcript per cell (Supplemental Fig. S5).

WTA signal is linearly correlated with RNAseq and RNAscope above a certain gene expression level, below which WTA does not detect signal (Fig. 2A and Supplemental Fig. S5). To identify this limit of quantitation of WTA relative to absolute transcript number, we performed breakpoint analysis, which fits two line segments to the data and iteratively calculates the breakpoint at which the model best fits the data. In 50 μm diameter AOIs containing an average of 13 cells, we found that the breakpoint was ~2 transcripts per cell. In larger AOI sizes with >50 cells, the breakpoint was 0.5-0.6 transcripts per cell (Supplemental Fig. S5), representing the lowest expression level that can be quantified by WTA.

Using genes with expression ≥1 expressed transcript per cell as measured by RNAscope as the true set of expressed genes, we calculated WTA sensitivity and specificity for targets with different absolute levels of gene expression. Specificity was high for all AOI sizes and ranged from 94-97% (Supplemental Fig. S5). Highly expressed targets (>10 transcripts per cell) were detected with a sensitivity of >80% in 50 μm diameter AOIs and 90-100% in larger AOIs. On the other extreme, very lowly expressed targets (1-2 transcripts per cell) were detected with a sensitivity of ~75% in AOIs with >500 cells and progressively less frequently detected in smaller AOIs (Fig. 2D). By combining the average number of transcripts per cell with the number of cells present in each AOI, we calculated sensitivity at different numbers of transcripts per AOI. At >100 transcripts per AOI, sensitivity was >70% (Fig. 2E). These results indicate that WTA can detect and quantify genes expressed at ~100 transcripts per AOI in AOIs ranging from 10-500 cells.

### WTA is compatible with multiple sample types and mouse strains

For the initial demonstration of the DSP technology, FFPE samples were used (Merritt et al. 2020). To expand the range of sample preparation types available for DSP, we designed and tested protocols for the use of WTA on human fresh frozen (FF) and mouse fixed frozen (FxF) samples (see Methods). To assess the performance of WTA on these additional sample types, we placed matched 200 μm diameter circular AOIs on FFPE and FxF mouse CPAs and FFPE and FF human tonsil tissue. The correlation of WTA counts was >0.8 comparing FFPE to either FxF or FF, and the distribution of signal to background ratios across genes was similar between sample preservation types (Supplemental Fig. S6). These results indicate that WTA results are concordant between FFPE and fixed frozen mouse and fresh frozen human tissues.

Specifically for mouse samples, we asked whether mouse WTA can accurately quantify gene expression in strains other than C57BL/6, which was used to generate the mouse reference transcriptome to which the panel was designed. To this end, we profiled an FFPE tissue array consisting of 7 different organs for each of 3 commonly used mouse strains (C57BL/6, BALB/c, and NOD/ShiLt) (Supplemental Fig. S7). Although transcriptional differences exist between strains due to true biological differences, these differences are known to be minimal (Breschi et al. 2017). We placed 300 μm diameter circular AOIs in similar regions of each tissue for each mouse strain and compared the results from each strain across organs. The transcriptomes were well correlated for all organs and pairs of strains, with correlation coefficients ranging from 0.7-0.95. Clustering by gene expression showed that organs clustered together before mouse strains, and gene expression patterns across tissues were similar in all three strains. These results suggest that despite small differences in the annotated transcriptomes, mouse WTA can be used to characterize gene expression in multiple strains.

### Whole transcriptome profiling of segmented regions reveals differences in spatial gene expression between tumor and tumor microenvironment across a range of AOI sizes

One of the strengths of the DSP system is that users can profile regions defined by morphology or expression of marker genes. The DSP instrument can segment a region of interest based on antibody- or RNA FISH-based fluorescence signals, splitting a single selected region into multiple AOIs (Fig. 3A). This feature enables different tissue compartments to be profiled separately even if they are spatially adjacent, as UV cleavage time has been optimized such that cross talk between regions is minimal (Merritt et al. 2020). We used this segmentation strategy to separate tumor and the tumor microenvironment (TME) to test whether WTA can detect expected differences in spatial gene expression in tissue. We also used this experiment as a model to assess the impacts of technical experimental design features such as AOI size and sequencing depth on WTA performance.

**Figure 3.**
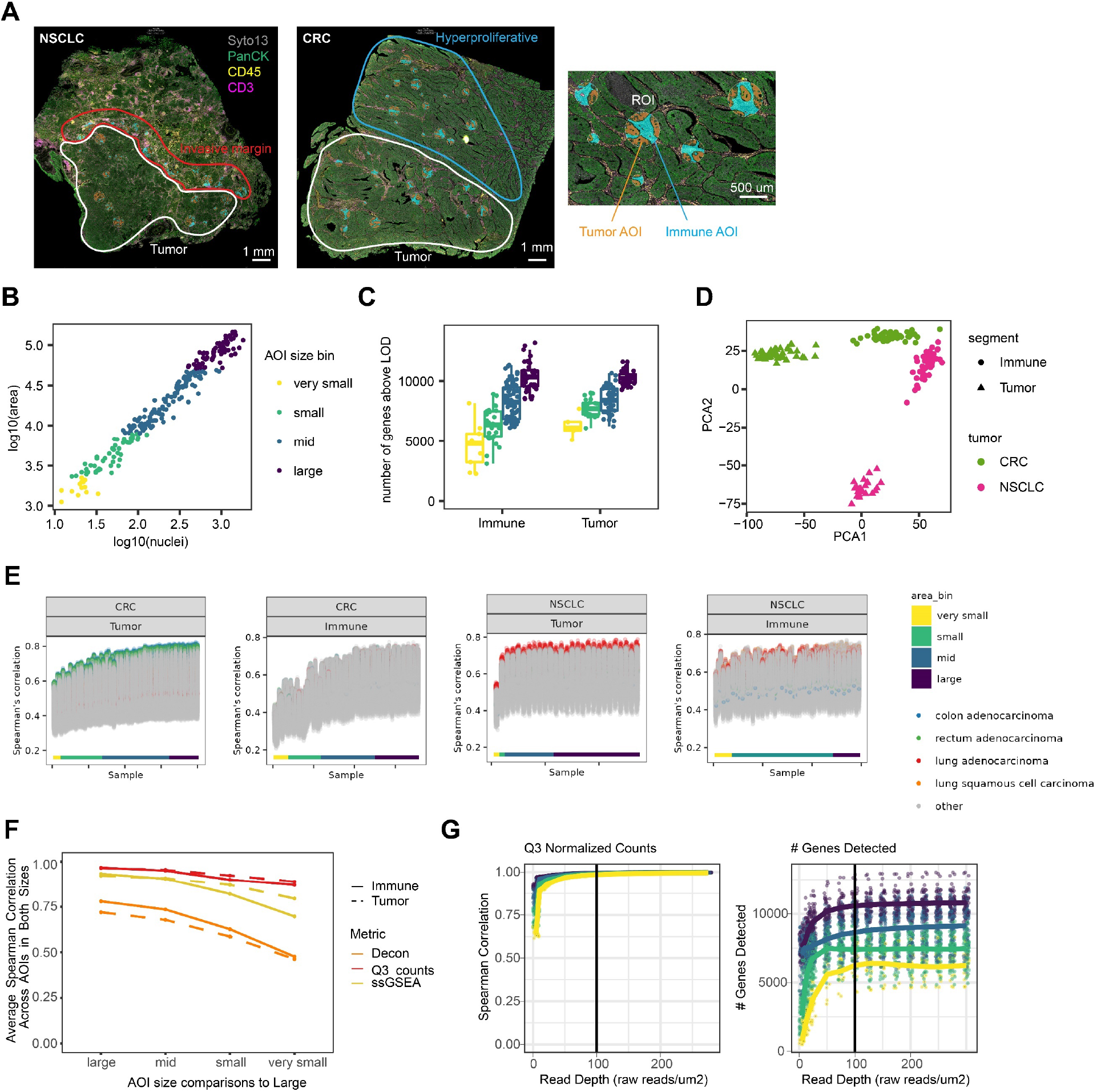
Effect of AOI size and sequencing depth on biological conclusions from segmented tumors and tumor microenvironment. **A.** Left: Representative images of the colorectal cancer (CRC) and non-small cell lung cancer (NSCLC) samples. Tumor, invasive margin, and hyperproliferative regions are highlighted. Slides were stained with antibodies against PanCK, CD3, and CD45. Right: Enlarged region of the CRC image to highlight the size titration and segmentation strategy. Circular regions of interest were automatically segmented into PanCK+ tumor (orange) and PanCK-immune (blue) compartments. **B.** Scatterplot of AOI area vs number of nuclei with points colored by the area bins used in the analyses in this figure. Very small: <2300 μm^2^, small: 2300-7850 μm^2^, mid: 7850-49,000 μm^2^, large: >49,000 μm^2^. **C.** Number of genes detected per AOI for tumor and immune compartments in each AOI size bin, colored as in B. **D.** Principal component analysis (PCA) of variation between samples using genes detected above background in >20% of AOIs. PCA1 vs PCA2 is plotted with points colored by tumor type and shaped by segment type. **E.** Spearman’s correlation of WTA counts from each AOI with all RNAseq datasets in the TCGA database. AOIs are ordered by area on the x-axis, and each point is a pairwise comparison with a dataset in TCGA. All genes in common between each pair of datasets were used in the correlation. Points are colored by tumor type in TCGA: colon adenocarcinoma (blue), rectal adenocarcinoma (green), lung adenocarcinoma (red), and lung squamous cell carcinoma (orange). All other tumor types are colored in grey. AOIs are labeled by area bin. **F.** Correlation of counts, single-sample Gene Set Enrichment Analysis (ssGSEA) enrichment, and cell type deconvolution between AOIs. For each of the three output metrics, Spearman’s correlations were calculated between each AOI, and averaged within different AOI size bins compared to the largest AOI sizes. AOIs are split into groups based on segment type (tumor or immune). **G.** Left: Spearman’s correlation of counts for each subsampled read depth and AOI size relative to counts at 300 reads/μm^2^. Right: Number of genes detected above background for each subsampled read depth and AOI size.

Two serial FFPE sections from colorectal cancer (CRC) and non-small cell lung cancer (NSCLC) samples were labeled with fluorescent antibodies against pan-cytokeratin (PanCK) to mark tumor, CD45 to mark immune cells broadly, and CD3 to mark T cells. After labeling the tissue with these morphology markers, we selected regions of interest in different pathological areas of the tissue: tumor and hyperproliferative regions in CRC samples and tumor and invasive margin regions in NSCLC samples. Regions were segmented by fluorescent antibody signal into tumor (PanCK+) and TME (PanCK-) AOIs (Fig. 3B).

To assess the effect of AOI size on WTA performance, we selected a range of circular region sizes and binned the resulting segmented AOIs into 4 size bins by area, a metric that is well correlated with cell count. Area bins ranged from “very small” (<2300 μm^2^ area, equivalent to a 55 μm diameter circle and with an average of 20 cells) to “large” (>49,000 μm^2^ area, equivalent to a 250 μm diameter circle and with an average of 920 cells) (Fig. 3B). WTA counts were well correlated between large and smaller AOIs: large AOIs had a median Pearson correlation of 0.94 with each other and very small AOIs had a median correlation of 0.71 with large AOIs (Fig. 3F, Supplemental Fig. S8). An increasing number of genes were detected above background in larger AOIs, with ~6000 genes detected per AOI in very small AOIs and ~11,000 genes detected in large AOIs (Fig. 3D). Genes detected in small AOIs were generally a subset of genes detected in large AOIs with very few genes detected only in small AOIs (Supplemental Fig. S8). Counts of the genes encoding the proteins used as markers for segmentation were highly enriched in the expected segment, and enrichments were similar for all AOI sizes (5-fold or greater median expression in the expected segment type) (Supplemental Fig. S8). We also confirmed that samples clustered by biological annotation (tumor type and tumor vs TME) regardless of AOI size (Fig. 3E).

We compared gene expression of each segmented AOI with all bulk RNAseq datasets in The Cancer Genome Atlas (TCGA) (Weinstein et al. 2013) to ask whether we could accurately classify the tumor type. We found that our classification of tumor segments was 100% accurate regardless of AOI size. All tumor segments correlated best with the expected tumor datasets in TCGA: colon adenocarcinoma and rectal adenocarcinoma for the CRC samples and lung adenocarcinoma for the NSCLC samples (Fig. 3E). Correlation coefficients increased with AOI size, from ~0.6 in very small AOIs to ~0.8 in large AOIs. As expected, TME segments generally did not correlate best with the matching tumor types in TCGA. The TME likely does not make up a substantial fraction of the tumors sequenced in the bulk TCGA datasets, highlighting the value of segmentation for capturing gene expression profiles of less abundant cell types.

We further examined whether we could detect the expected biological differences between tumor and TME in AOIs of different sizes. Immune-related pathways, such as interleukin signaling and tumor necrosis factor signaling, were enriched in TME, while pathways related to cell motility, proliferation, and cancer-associated signaling were enriched in tumors (Supplemental Fig. S8). Pathway analysis results were well correlated between AOIs of the same type and between large and small AOIs (Fig 3F). We next performed cell type deconvolution with SpatialDecon, an algorithm for estimating abundance of cell types defined by single cell sequencing in spatial gene expression data (Danaher et al. 2022). We used gene expression profiles of immune and stroma cells to characterize the immune cell content of tumor and TME segments. As expected, tumor segments had a very low estimated abundance of immune cells relative to TME for all AOI sizes (Supplemental Fig. S8), highlighting the ability of segmentation to separate cell types. Cell type deconvolution results were more variable between individual AOIs in all size bins, but correlation decreased with size to a similar degree as other metrics (Fig. 3F). These results demonstrate the robustness of the WTA for biological characterization across a wide range of AOI sizes.

In addition to AOI size, we also assessed the impact of sequencing depth on WTA data. All AOIs were deeply sequenced and reads were subsampled in silico from a read depth of 5 raw reads/μm^2^ to 300 raw reads/μm^2^. Five replicates of the subsampling were performed at each read depth. As expected, increasing read depth corresponds to a lower fraction of unique UMIs, indicating higher sequencing saturation of the libraries (Supplemental Fig. S9). Small AOIs reached higher saturation at lower read depths than larger AOIs, consistent with lower starting molecular complexity in these samples. For each subsampled dataset, we compared the number of genes detected and correlations of counts, pathway enrichment results, cell type deconvolution results, and differential expression results to the highest sequencing depth. For most metrics and AOI sizes, results were well correlated at all but the lowest sequencing depths and stabilized by 100 raw reads/μm^2^, corresponding to a sequencing saturation of ~50%. Correlation of cell type deconvolution results did continue to improve with higher read depth, especially in small AOIs, suggesting that robust deconvolution might benefit from higher sequencing saturation (Supplemental Fig. S9).

### Profiling transcriptomes of anatomical structures in normal kidney and kidney disease

To demonstrate the capability of WTA to integrate the transcriptome with annotated histological and pathological features of a tissue, we asked how the transcriptome is altered in anatomically distinct regions of the kidney with diabetic kidney disease (DKD). The kidney nephron has a complex structure that includes the glomerulus, a cluster of specialized cells that forms the filtration barrier, and the tubule, which reabsorbs water and small molecules and has different functions along its length. The effects of DKD on the glomeruli have been well studied, such as a loss of glomerular filtration, inflammation, and immune cell infiltration (Reidy et al. 2014; Thomas et al. 2015). However, DKD affects all parts of the kidney. Therefore, we used WTA to profile the transcriptome of three nephron substructures: the glomeruli, the proximal convoluted tubules, and the distal convoluted tubules.

We profiled three normal and four DKD FFPE human kidney samples. To identify and discriminate kidney structures, three fluorescently labeled antibodies targeting epithelia (PanCK), immune cells (CD45), and podocytes (WT1) in glomeruli were used. Glomeruli and tubules were identified morphologically and polygon-shaped AOIs were drawn to capture each structure. Tubules were segmented based on the PanCK signal into proximal (PanCK-) and distal tubules/collecting duct (PanCK+) (Fig. 4A). Within each sample, individual glomeruli were annotated by a pathologist for severity of disease-related changes using both the fluorescence images and hematoxylin and eosin (H&E) images of serial sections. The data were collected from both relatively healthy and more abnormal glomeruli in both normal and DKD samples (Fig. 4B).

**Figure 4.**
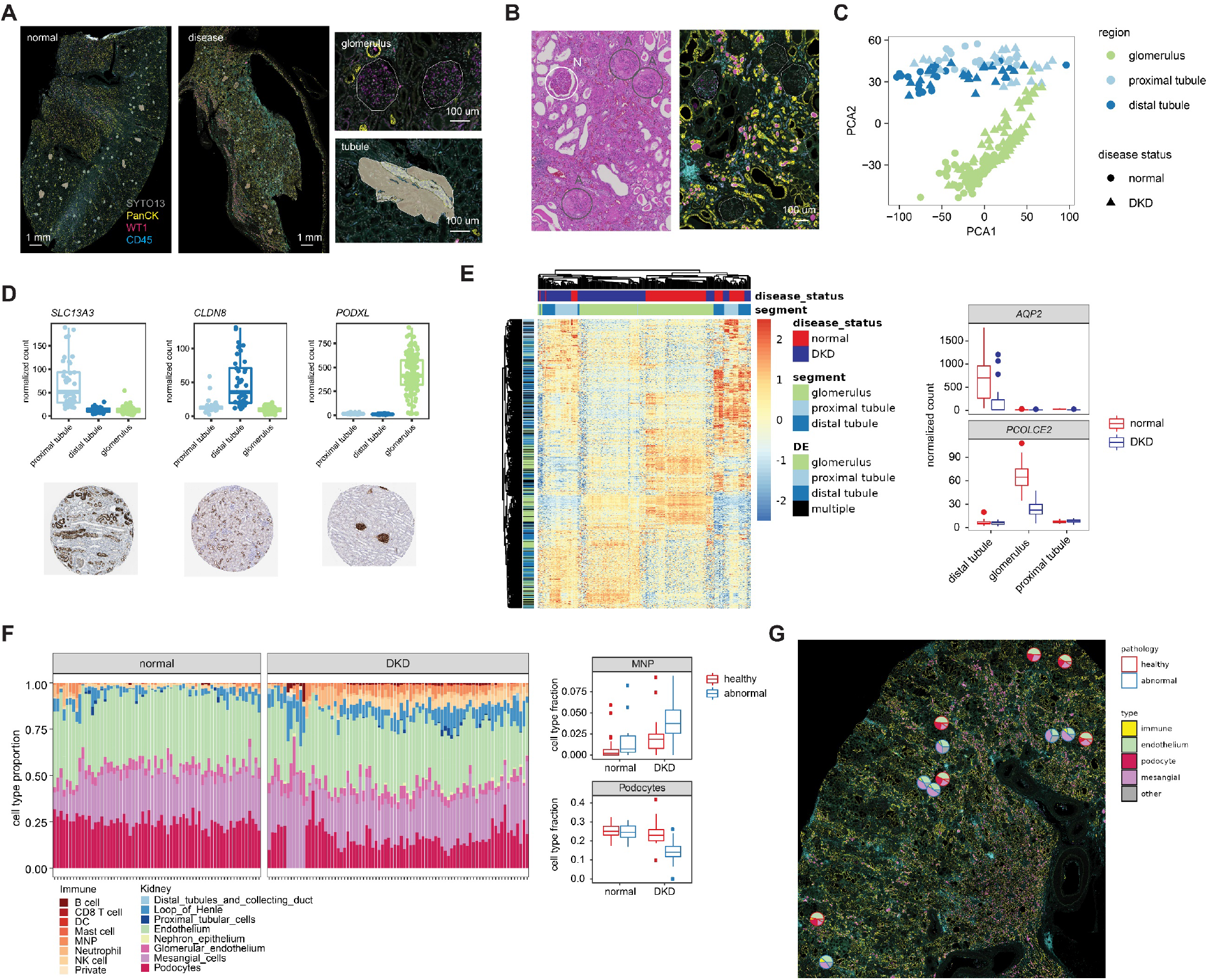
Spatial heterogeneity in gene expression changes associated with diabetic kidney disease in human kidneys. **A.** Left: Representative fluorescence images of normal and diabetic human kidneys. Tissues were stained with antibodies against PanCK, WT1, and CD45. Right: Example images from normal kidney highlighting the AOI strategy. Glomeruli were profiled using polygon-shaped AOIs, and tubules were automatically segmented in proximal tubules (PanCK-) and distal tubules (PanCK+). **B.** Individual glomeruli in each kidney sample were annotated by degree of pathology. A representative H&E image (left) and fluorescence image (right) from the same region of a diabetic kidney specimen are shown. Glomeruli with higher degree of abnormality are circled in grey and labeled “A”, while those that are more normal are circled in white and labeled “N”. **C.** Principal component analysis of variation between samples using genes detected above background in >1% of AOIs. PCA1 vs PCA2 is plotted, with substructure indicated by color and disease status indicated by shape. **D.** Boxplots of counts in all AOIs of three example genes differentially expressed between kidney substructures with corresponding antibody staining images from the Human Protein Atlas (https://www.proteinatlas.org/) (Uhlén et al. 2015). **E.** Left: Heatmap of most differentially expressed genes between normal and DKD in glomeruli, distal tubules, and proximal tubules. All genes are significant at FDR <0.05 and a fold change of >1.5. Genes are annotated by the structure in which they were significantly differentially expressed, or “multiple” for the genes significant in more than one structure. Columns and rows are clustered by hierarchical clustering and the data are scaled by row. Right: Boxplot of normalized counts for two example genes in normal and DKD glomeruli, proximal tubules, and distal tubules. **F.** Left: Results of cell type deconvolution of glomeruli using single-cell expression data from (Young et al. 2018). Data are displayed as stacked barplots with each bar as a single AOI and the estimated proportion of each cell type colored, and faceted by disease status. Right: Boxplots of proportions of two example cell types with significantly different proportions in normal and DKD glomeruli (t-test Bonferroni-corrected p-value <0.05), colored by whether the glomerulus was annotated as pathologically abnormal or healthy. MNP = mononuclear phagocyte, DC = dendritic cells. **G.** Pie charts overlaid over the fluorescence image of a single kidney showing the proportion of different glomerulus and immune cell types for each glomerulus profiled in a representative disease sample. Each plot is outlined based on pathological annotation: abnormal glomeruli (blue), healthy glomeruli (red).

Overall, we profiled 231 AOIs that passed quality filters, across which we detected and quantified 16,084 genes. AOIs clustered by region and by disease status more closely than by patient (Fig. 4C, Supplemental Fig. S10). In normal kidneys, we identified over 6000 significantly differentially expressed genes between glomeruli and tubules, and over 8000 differentially expressed genes between proximal and distal tubules. We found a strong concordance between genes differentially expressed in our study and those differentially expressed between cell types in kidney single-cell RNAseq (Young et al. 2018) (Supplemental Fig. S10). Furthermore, we validated example genes differentially expressed in each structure with publicly available antibody staining from the Human Protein Atlas (Uhlén et al. 2015), and saw excellent concordance of spatial localization (Fig. 4D).

At the pathway level, differentially expressed pathways between glomeruli, proximal, and distal tubules recapitulated known aspects of kidney biology. For example, pathways specifically enriched in proximal tubules included anion and amino acid transporters, which are known to be highly expressed in proximal tubules, while bicarbonate transporters were enriched in both proximal and distal tubules. Pathways enriched in glomeruli include nephrin and SEMA3A signaling, which are key proteins expressed in cells of the glomerular filtration membrane (Reidy and Tufro 2011; Martin and Jones 2018) (Supplemental Fig. S10).

With DKD, we observed 2400 differentially expressed genes across the different kidney substructures compared to normal kidney samples. For most genes dysregulated with disease, expression changes were correlated across the different anatomical structures, but some structure-specific genes were altered with disease (Fig. 4E). For example, the gene *PCOLCE2* is only expressed in glomeruli and is substantially downregulated with disease. Expression of this gene has been observed in glomerular podocytes, a specialized cell that forms the glomerular filtration barrier, and lower expression correlates with loss of renal function in chronic kidney disease patients (Ju et al. 2013). Similarly, aquaporin genes such as *AQP2* and *AQP3* are strongly downregulated in the distal tubules with disease. This family of genes encodes water channels necessary for concentration of urine by the kidneys and is specifically expressed in tubules (Nielsen et al. 1999). These results indicate that DKD can cause loss of substructure-specific and cell type-specific gene expression critical for normal kidney function.

Loss of glomerular podocytes and increased immune cell infiltration are known to be hallmarks of DKD. We recapitulated this phenotype using cell type deconvolution with the SpatialDecon algorithm using gene expression signatures from published kidney single cell RNA sequencing data (Young et al. 2018). We observed a marked loss of podocytes in glomeruli and increased abundance in almost all types of immune cells in all substructures (Fig. 4F, Supplemental Fig. S10). Interestingly, we identified that the loss of podocytes was heterogeneous across individual glomeruli. Even within diseased or normal samples, pathologically abnormal glomeruli had a more profound loss of podocytes and higher levels of immune infiltration compared to glomeruli with fewer pathological features. In particular, the abundance of B cells, natural killer cells, and mononuclear phagocytes increased in diseased kidneys but the increase was significantly higher in more severely pathologically abnormal glomeruli (Fig. 4F, Supplemental Fig. S10). This spatial heterogeneity was observed within individual diseased kidneys (Fig. 4G), indicating that some glomeruli are more affected by disease despite close physical proximity. In total, these results demonstrate the feasibility of whole transcriptome profiling of specific organ substructures to detect spatially variable disease-related abnormalities.

### Identifying organ substructure-specific transcriptomes in the developing mouse embryo

One anticipated use of WTA is to catalog spatial gene expression profiles in histological structures and anatomical regions across organs. To demonstrate the utility of WTA for building spatial organ atlases, we profiled whole transcriptomes of different organs and organ substructures in a developing mouse embryo. A single fixed-frozen E13.5 mouse embryo was sectioned along the sagittal plane. Six sections spanning the embryo were stained with antibodies against TRP63 (epithelial marker) and β3 tubulin (neuronal microtubule marker) and hybridized with mouse WTA probes (Fig. 5A).

**Figure 5.**
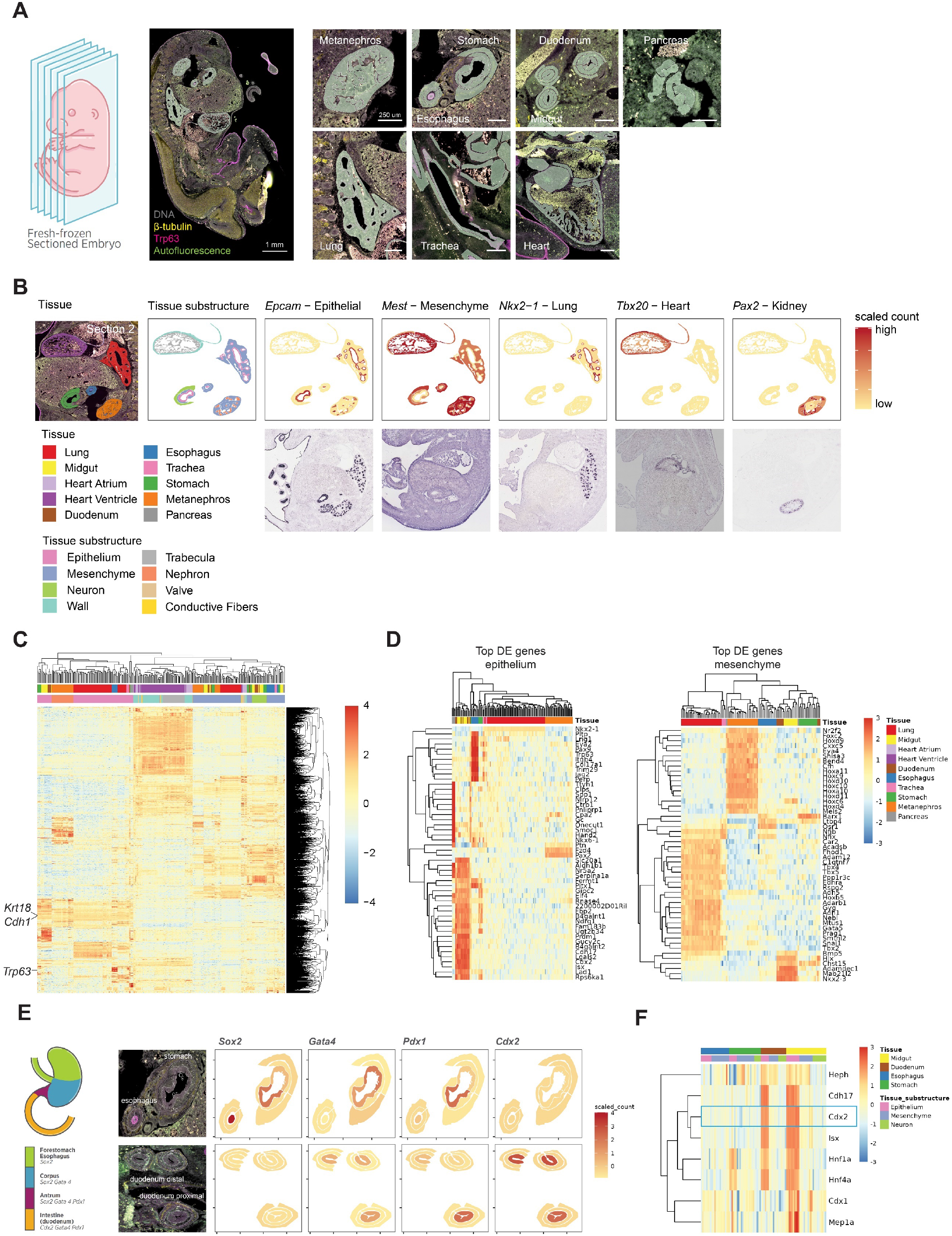
Spatial profiling of transcriptional programs during organogenesis in a mid-gestation mouse embryo. **A.** Left: Schematic and representative image of the fixed-frozen E13.5 mouse embryo profiled. Sections were labeled with antibodies against TRP63 (magenta) and β3 tubulin (yellow). Autofluorescence is shown in green. Right: Example images of each organ profiled showing the AOI profiling strategy. Freeform polygon AOIs capture anatomical substructures of each organ. **B.** Expression of marker genes for specific organs and cell types in an example section compared to ISH images of the same genes in E14.5 mouse embryos from the GenePaint database (https://gp3.mpg.de/, Diez-Roux et al. 2011; Visel et al. 2004). Tissue, tissue substructure, or normalized scaled WTA count for five genes is plotted over the shape of each AOI. **C.** Heatmap showing scaled expression of the 2000 most variable genes across the dataset. Columns and rows are clustered by hierarchical clustering and columns are annotated by organ and organ substructure. **D.** Heatmaps showing scaled expression of the top 50 most differentially expressed genes in epithelium (left) and mesenchyme (right). All genes shown are significant at Bonferroni-corrected p-value < 0.01. Columns and rows are clustered by hierarchical clustering and columns annotated by organ. **E.** Left: Schematic of key transcription factor expression in stomach and gut development (adapted from (Willet and Mills 2016)). Right: Expression of the same transcription factors plotted on example AOIs from a representative section. **F.** Heatmap showing scaled expression of *Cdx2* and *Cdx2* target genes from (Gao et al. 2009) in esophagus, stomach, duodenum, and midgut AOIs. Columns and rows are clustered by hierarchical clustering and columns are annotated by organ and organ substructure.

AOIs were selected in 9 organs across the 6 sections (heart, lung, metanephros, pancreas, midgut, duodenum, stomach, esophagus, and trachea). Within each organ, freeform polygon-shaped AOIs were drawn to capture morphologically distinct substructures using anatomical features identified using both the fluorescence image and an H&E-stained serial section (Fig. 5B, Supplemental Fig. S11). For example, in the developing heart, we placed AOIs in the ventricle wall, atrium wall, trabeculae, conductive fibers, and valves. In the stomach, esophagus, duodenum, and midgut, we selected AOIs in the epithelial, neural, and mesenchymal layers.

Overall, we profiled the whole transcriptome of 347 AOIs across the nine organs and 2-5 substructures per organ. We identified 17,662 genes expressed above background, indicating that in diverse tissues nearly the entire transcriptome is detectable by WTA. Examining the spatial expression of cell-type-specific marker genes showed the expected patterns; for example, the epithelial marker *Epcam* is expressed in epithelial AOIs in all tissues while the mesenchymal marker *Mest* is highly expressed in the mesenchyme and heart but not in the epithelial AOIs. We observed that genes known to be highly expressed in specific tissues were restricted to the expected tissue, but still show spatial variability within a tissue. For example, the lung transcription factor *Nkx2-1* was expressed in the lung and trachea epithelium, as has been previously reported (Minoo et al. 1999), and the kidney transcription factor *Pax2* was specifically expressed in AOIs in the metanephros cortex (Bouchard et al. 2002; Minoo et al. 1999). The spatial expression pattern of these tissue or substructure-specific genes matched the observed expression pattern by colorimetric RNA ISH of E14.5 mouse embryos (Visel et al. 2004; Diez-Roux et al. 2011), validating the WTA results (Fig. 5B).

Clustering AOIs by gene expression reveals that heart AOIs cluster separately from the other organs, and that for the non-heart AOIs similar substructures cluster together first, and then by organ. Epithelial AOIs form one cluster, as do mesenchymal and neuron AOIs. Within each substructure, both common and tissue-specific genes can be identified. Across the epithelial AOIs, shared highly expressed genes include known epithelial markers *Cdh1* and *Krt18*. Among tissue-specific epithelial genes, *Trp63* was expressed only in the epithelium of the esophagus and trachea, matching the expression pattern of the TRP63 antibody morphology marker used in this study (Fig. 5C).

As most of the organs profiled have an epithelial and mesenchymal region, we identified genes differentially expressed between organs in these two cell types (Fig. 5D). Organ-specific genes were nearly non-overlapping between epithelium and mesenchyme, highlighting the value of capturing substructure-specific transcriptomes over bulk organ gene expression profiling. Amongst the top organ-specific genes include key developmental transcription factors: *Nkx6-1,* a critical regulator of pancreas β cell development (Aigha and Abdelalim 2020), was uniquely expressed in the pancreas epithelium; *Cdx2,* an intestine-specific transcription factor necessary for intestine differentiation (Gao et al. 2009), was expressed in the duodenum and midgut epithelium; and *Barx1,* which is necessary for stomach differentiation (Kim et al. 2005), was localized to the stomach mesenchyme.

As developmental transcription factors were among the most differentially expressed genes across organs and organ substructures, we next asked whether our data could recapitulate the known developmental specification of the digestive system in mid-gestation embryos. Around E13, the transcription factors *Sox2, Gata4, Pdx1,* and *Cdx2* are localized in an overlapping pattern from anterior to posterior in the developing esophagus, stomach, and intestine and are necessary for proper specification of those tissues. For example, *Sox2* is expressed in the developing esophagus and stomach, while *Cdx2* is expressed in the intestine. Loss of *Cdx2* in the intestine leads to the misexpression of *Sox2* in that tissue and the ectopic expression of stomach and esophageal markers (Kumar et al. 2019; Willet and Mills 2016). Our data accurately recapitulated this known pattern of transcription factor expression across tissues and revealed spatial patterns within each tissue (Fig. 5E). All four transcription factors were predominantly located to the epithelium in each tissue, and *Pdx1* is more highly expressed in the liver proximal section of the duodenum than the distal section. Furthermore, we examined the expression of pro-intestinal targets of *Cdx1* in digestive system AOIs (Raghoebir et al. 2012). Several canonical *Cdx2* targets, such as *Cdh17,* were expressed in the same spatial pattern as *Cdx1,* which is limited to the intestinal epithelium. However, others were expressed more broadly or more narrowly, such as *Hnf1a* and *Hnf4a,* which were also expressed in stomach epithelium, and *Heph,* which was also expressed in the intestinal mesenchyme, suggesting more complex regulation governing the expression of these genes (Fig. 5F). Overall, these results demonstrate the capacity of WTA to reveal the complex spatial gene expression patterns governing key cell fate decisions during embryonic development.

## Discussion

The Whole Transcriptome Atlas is a high-plex in situ hybridization method for spatial transcriptome profiling using the Digital Spatial Profiling platform. Here we describe the design, performance, and applications of the human and mouse WTAs, which comprise >18,000 multiplexed probes targeting the protein-coding genes of the human or mouse transcriptome. We show that WTA data is reproducible and concordant with orthogonal gene expression profiling methods and can quantify genes with low, medium, and high expression levels depending on the size of the profiled region. Furthermore, the applications of WTA to human disease biology and mouse developmental biology demonstrate that whole transcriptome data enables comprehensive pathway-level spatial analyses.

DSP technology allows flexible and customizable region selection that can trace the boundaries of anatomical or biological structures or groups of specific cells. As a result, a wide range of AOI sizes and types are possible, from a minimum region size of 5 μm x 5 μm to a maximum of 660 μm × 785 μm (Bergholtz et al. 2021). Smaller regions have the advantage of less heterogeneity and higher spatial resolution, but as with other existing spatial gene expression methods profiling smaller regions results in lower sensitivity. To define these tradeoffs, we benchmarked the sensitivity of WTA using differently sized AOIs in homogeneous cell pellets, which have the advantage of not being confounded by spatial variation such that data can be directly compared with bulk RNAseq. We find that in AOIs with ~100 cells, we detect ~70% of the genes observed in bulk RNAseq, a high sensitivity given that that bulk RNAseq is based on tens to hundreds of thousands of cells as input. Using RNAscope, we demonstrate that WTA sensitivity is equivalent to <1 transcript/cell in AOIs of at least 100 cells, with ~100 transcripts required per AOI for robust detection. In tumor tissue, this sensitivity corresponds to detecting ~6000 genes in small AOIs with <20 cells, and >10,000 genes in large AOIs with hundreds of cells. As expected, there is a tradeoff between WTA signal and AOI size: more genes detected above background, better coverage of low expressing genes, and higher reproducibility in larger AOIs. However, we demonstrate that WTA counts from small AOIs still correlate well with orthogonal gene expression methods, and that the results of downstream analyses such as clustering, differential expression, and pathway enrichment are relatively robust to AOI size. These findings will enable researchers to devise a profiling strategy that is suited to address their specific experimental question.

Transcriptome-scale spatial data enables a wide range of pathway-level downstream analyses. With WTA, we detect expected pathways enrichment in the glomeruli and tubules of human kidneys, and also demonstrate robust detection and spatial localization of the key transcription factors and their target genes in mouse organogenesis. Furthermore, methods such as cell type deconvolution allow the integration of gene expression signatures from single-cell RNAseq data with spatial data, enabling the localization of specific cell types in space (Danaher et al. 2022). In this work, we demonstrate heterogeneity in cell type loss in diabetic kidney disease that can be linked to pathological annotation of the tissue. The integration of scRNAseq and WTA spatial analysis has been demonstrated in other contexts as well, including in pancreatic ductal adenocarcinoma to reveal that a malignant cell type identified by scRNAseq was spatially associated with higher immune infiltrations (Hwang et al.).

The development of a whole transcriptome panel for both human and mouse enables a wide range of translational, clinical, and basic biology research. For example, researchers have used WTA to identify focal changes in gene expression in kidney allograft rejection (Salem et al. 2022), determine cell types affected by SARS-CoV-2 infection in the olfactory epithelium (Khan et al. 2021), characterize functional gene expression differences across a heterogenous central nervous system tumor (Dottermusch et al. 2021), and assess structure-specific responses to treatment in prostate hyperplasia (Joseph et al. 2022). To promote these broad research applications, we have shown that WTA is compatible with a variety of tissue types and sample preservation methods (FFPE, fresh frozen, and fixed frozen). Moreover, we demonstrate successful WTA experiments in diverse normal and diseased human tissue, and in a wide range of tissues in adult and developing mice. One limitation of an ISH-based technology is that new probes must be designed to target each transcriptome of interest. However, we show that mouse WTA is compatible with the most commonly used mouse strains despite small differences in transcript sequence. In addition, WTA can be supplemented with custom-designed probes targeting additional transcripts of interest. For example, Delorey et al. used human WTA with an additional 26 probes designed against SARS-CoV-2 transcripts to create a spatial atlas of gene expression in different anatomical substructures and levels of virus infection in COVID-19 infected lungs (Delorey et al. 2021). Similarly, custom probes can be added to WTA to quantitate different transcript isoforms and non-coding RNAs, as WTA does not use poly-adenylated transcript capture.

Spatial gene and protein expression profiling with DSP has enabled discoveries in many research fields including oncology, immunology, neuroscience, and infectious disease. WTA expands the capabilities of DSP RNA profiling from 1,400 genes to the whole transcriptome level and enables high-plex spatial profiling of both human and mouse tissues. Future research will combine spatial whole transcriptome profiling with complex annotations and with sample timepoints to provide high-dimensional profiles of development, disease progression, and other biological processes.

## Methods

### Design of the Whole Transcriptome Atlas probes

The NCBI RefSeq reference transcriptomes for human (GRCh38.p13) and mouse (GRCm38.p6, C57BL/6) were used for design of human and mouse WTA, respectively. The genes targeted for design included all protein-coding genes with a few exceptions. Human protein-coding genes were determined based on the HUGO Gene Nomenclature Committee (HGNC) and designed according to the available RefSeq transcripts. Mouse protein-coding genes were determined based on Mouse Genome Informatics (MGI) and designed according to the available RefSeq transcripts. For mouse genes, we also considered the current status of genes in NCBI RefSeq and did not include those with poor status (Suppressed, Provisional, Model, or Inferred). Notably, 1450 protein-coding genes that exist in the MGI database had no available mRNA transcripts in RefSeq at the time of design. By comparison, that number in human was only 31 and included a few genes that should have been characterized as loci and not protein-coding entities (ex. PCDHG@, TRD).

In order to provide the best sensitivity for lower-expressing transcripts, we elected to remove the top 10 most highly expressed genes in TCGA across tumor types from the human WTA *(ACTB, ACTG1, EEF1A1, EEF2, FTL, GAPDH, PSAP, RPL3, TPT1,* and *UBC).* A similar assessment was performed for mouse genes according to (Söllner et al. 2017) but as most of the genes identified were organ-specific, we opted to instead remove genes based on empirical data using our assay. Those genes were *Gm20594* and *Eef1a1. Eef1a1* is the mouse homolog of human *EEF1A1* we prospectively removed for the same rationale, and *Gm20594* is the human ortholog of *MTRNR2L7,* which has homology to mitochondrial rRNA and thus could yield very high counts. In both human and mouse WTA, mitochondrially-encoded transcripts were removed as they are also very highly expressed.

The probe design process begins with an exhaustive evaluation of all possible contiguous 35-50 nucleotide sequence windows for each mRNA target. This large pool of possible probe candidates is first filtered for intrinsic characteristics including melting temperature, GC content, secondary structure, and runs of poly-nucleotides. Probes satisfying these parameters are further screened for homology to the full transcriptome of the parent organism utilizing the Basic Local Alignment Search Tool (BLAST) from the National Center for Biotechnology Information (NCBI). Preference was given to probes with absence of homology to off-target genes in the corresponding transcriptome, covering known protein coding transcripts, lying within coding regions, and maximizing the coverage of the isoform repertoire. Targeting of a transcript was judged based on ≥95% sequence identity to the probe target. Previous work has found that selecting probes that are 95-100% identical to the intended target and filtering out probes that are ≥75-85% in homology and that possess ≥15-17 MCB (Maximum Contiguous Bases) confer excellent specificity to the intended target (Militon et al. 2007; Kane 2000; Rimour et al. 2005). Final panel candidates are further screened for intermolecular interactions with other probes in the candidate pool including potential probe-probe hybridization as well as minimizing common sequences between probes.

For both human and mouse WTA, negative control probes were designed against synthetic sequences from the External RNA Controls Consortium (ERCC) set (Baker et al. 2005). Negative control probes were designed to have similar GC and T_m_ properties as target probes and are subject to the same intermolecular interaction screening. The final probe pool consists of 18,815 probes for human WTA and 20,175 probes for mouse WTA, including 139 negative control probes for human and 210 negative control probes for mouse. These probes target 19,505 and 21,596 annotated genes for human and mouse, of which 19,128 and 21,040 are protein-coding respectively. Due to high homology in some gene families, 636 human probes and 656 mouse probes target more than one gene (Supplemental Table S1).

Probes contain an indexing sequence separated from the RNA-targeting region by a UV-photocleavable linker (Supplemental Fig. S1). The indexing sequence contains a 12-nucleotide barcode identifying the RNA-targeting sequence, a 14-nucleotide random UMI, and primer binding sites for the amplification of tags and addition of P5 and P7 adaptors for Illumina NGS. The RNA ID barcodes were designed to have a minimum Hamming distance of ≥2 between barcodes.

For the RNA FISH comparison experiments and the CRC and NSCLC experiments, an early version of the human WTA probe pool was used that differed slightly from the final commercially available version used for all other experiments. For these experiments, probes were filtered to only include those in the commercially available pool before any analyses were performed.

### Sample preparation for DSP

Sample preparation was performed as described in the NanoString GeoMx RNA-NGS slide preparation manuals. Maximum sample size for imaging on the DSP instrument is 36.2 mm long by 14.6 mm wide (Bergholtz et al. 2021). Samples were processed on a Leica Bond RX or RXm automated stainer (Leica Biosystems) or manually. For FFPE samples, freshly cut 5 μm sections were mounted on positively charged slides, baked, deparaffinized, washed in ethanol, and washed in PBS or Leica Bond Wash Solution. Targets were retrieved in Tris-EDTA pH 9.0 in a pressure cooker (manual protocol) or Leica BOND Epitope Retrieval Solution (automated protocol) for 10 min at 85°C (cell pellets), 10 min at 100°C (tonsil), or 20 min at 100°C (human CRC, human NSCLC, human kidney, and mouse tissue arrays), and washed in PBS or Bond Wash Solution. Samples were digested with 0.1 mg/mL proteinase K for 5 min (cell pellets) or 1 μg/mL for 15 min (tissues) at 37°C and washed with PBS. For fresh frozen human tonsil samples, 5 μm sections were mounted on positively charged slides and fixed overnight in 10% NBF. Antigen retrieval, digestion, and washes were performed as described for FFPE except that the proteinase K digestion was at room temperature. Fixed frozen mouse cell pellets (Acepix Biosciences) were fixed in 4% PFA overnight at 4°C, embedded in OCT, and snap frozen. Fixed frozen mouse embryos (Acepix Biosciences) were fixed in 10% NBF overnight at room temperature, embedded in OCT, and snap frozen. For both cell pellets and embryos, 10 μm OCT embedded sections were washed in PBS, washed in ethanol, and antigen retrieval was performed for 15 min at 85°C (embryo) or 10 min at 85°C (cell pellets). Digestion and washes were performed as for FFPE.

All samples were incubated overnight at 37°C with human or mouse WTA following the NanoString GeoMx RNA-NGS slide preparation manual at a probe concentration of 4 nM per probe in 2x SSC with 2.5% dextran sulfate, 0.2% BSA, 100 μg/mL salmon sperm DNA, and 40% formamide. During incubation, slides were covered with HybriSlip Hybridization Covers (Grace BioLabs). After incubation, coverslips were removed by soaking in 2x SSC + 0.1% Tween-20. Two 25 min stringent washes were performed in 50% formamide in 2x SSC at 37°C to remove unbound probe, and samples were washed in 2x SSC. For antibody morphology marker staining, samples were incubated in blocking buffer for 30 min at room temperature in a humidity chamber, and then incubated with 100 μm SYTO13 and the relevant fluorescently conjugated antibodies (Supplemental Table S2) for 1-2 hours. Samples were washed in 2x SSC and loaded on the GeoMx DSP instrument.

### Fluorescent in situ hybridization with RNAscope

ISH was performed using the RNAscope LS Multiplex Fluorescent Reagent kit (ACD) using a Leica Bond RX or RXm automated stainer according to the manufacturer’s instructions. Antigen retrieval was performed for 15 min at 88°C, and digestions were performed with ACD protease for 15 min at 40°C. A list of probes used is in Supplemental Table S2. Probes were visualized with TSA plus Cy3, Cy5, or Opal620.

RNAscope spot counting was performed as previously described (Merritt et al. 2020). Briefly, slides were imaged using the Nikon Eclipse TE2000-E microscope at 40x magnification. Images were captured with Nikon Elements commercial software. For imaging, *z* stacks at 0.5 μm steps were taken from the top to bottom focal planes of each cell pellet. Exposure time was set manually to have maximal signal for the lowest expressing cell line while remaining non-saturated for the highest expressing cell line. Maximum *z*-projection images were created with Nikon Elements software across all channels. QuPath software (https://qupath.github.io/) was used to quantify the number of RNAscope spots and cells imaged per field of view using the method and scripts described in (Merritt et al. 2020).

For the comparison of total RNAscope fluorescence intensity with WTA counts, the mean pixel intensity of each AOI for each relevant channel in the DSP 20x scan image was extracted and multiplied by total AOI area to get total fluorescence intensity.

### DSP experiments

DSP experiments were performed according to the NanoString GeoMx-NGS DSP Instrument manual and as previously described (Merritt et al. 2020; Bergholtz et al. 2021). Briefly, slides were imaged in four fluorescence channels (FITC/525 nm, Cy3/568 nm, Texas Red/615 nm, Cy5/666 nm) to visualize morphology markers, and regions of interest were selected for collection. For the CRC/NSCLC and the kidney experiment, regions of interest were segmented based on the expression of morphology markers using the DSP auto-segmentation tool with manually tuned settings. AOIs were illuminated and released tags were collected into 96-well plates as previously described.

### Next-generation sequencing and sequencing data analysis

Library preparation for NGS was performed according to the NanoString GeoMx-NGS Readout Library Prep manual. Briefly, the DSP aspirate was dried and resuspended in 10 uL DEPC-treated water, and 4 μL were used in a PCR reaction. NanoString SeqCode primers were used to amplify the tags and add Illumina adaptor sequences and sample demultiplexing barcodes. PCR products were pooled either in equal volumes or in proportion relative to AOI size, depending on the experiment, and purified with 2 rounds of AMPure XP beads (Beckman Coulter). Libraries were sequenced on an Illumina NextSeq 550, NextSeq 2000, or NovaSeq 6000 according to the manufacturer’s instructions, with at least 27×27 paired end reads.

FASTQ files were processed using the NanoString GeoMx NGS Pipeline v2.0 or v2.2. Briefly, reads were trimmed to remove low quality bases and adapter sequences. Paired end reads were aligned and stitched, and the barcode and UMI sequences were extracted. Barcodes were matched to known probe barcodes with maximum 1 mismatch allowed. Reads matching the same barcode were deduplicated by UMI. The number of raw reads was highly linearly correlated with the number of unique UMIs at all AOI sizes, suggesting largely uniform library amplification (Supplemental Fig S9). However, to correct for any PCR amplification bias, all analyses in this study use UMI deduplicated counts.

### RNAseq experiments

For the comparison to cell line RNAseq, in-house RNAseq data was generated for all of the mouse cell lines used in the comparison and 5 of 11 human cell lines (Daudi, H596, HEL, HUT78, and HS578T). Purified total RNA for each cell line was purchased from Acepix Biosciences. RNAseq libraries were prepared using the TruSeq Stranded mRNA Library Prep kit (Illumina) following the manufacturer’s instructions and using 100-125 ng of RNA per cell line as input. Libraries were sequenced on an Illumina NextSeq 550 with 75×75 paired end reads.

Sequencing reads were mapped to the human RefSeq transcriptome GRCh38.p13 or the mouse reference transcriptome GRCm38.p6 using Salmon v1.3.0 with default parameters (Patro et al. 2017). Transcript-level counts were collapsed to gene-level counts using tximport v3.13 (Soneson et al. 2015).

Comparison of our in-house human cell line RNAseq data to publicly available RNAseq data from the Cancer Cell Line Encyclopedia Project (CCLE) (Ghandi et al. 2019) showed that our data was highly correlated with the CCLE data, and that WTA correlations and sensitivity were very similar using our data and the CCLE data. As there was CCLE RNAseq data available for all of the human cell lines profiled by WTA, the CCLE data was used for all of the comparisons to human WTA shown in Figure 1 and Figure S2.

### Data analysis and visualization

Count data was processed and normalized using either the NanoString DSPDA software v2.2 or v2.3, the GeoMxTools R package v1.0 (https://bioconductor.org/packages/release/bioc/html/GeomxTools.html), or similar in house data processing scripts. AOIs with fewer than 5000 raw reads or a sequencing saturation <45% (mouse embryo experiment) or <50% (all other experiments) were filtered out of the analysis. For the negative probes, we performed outlier testing and removed outlier probes from the analysis before collapsing counts. All other targets have just one probe per target and therefore were not filtered for outliers or collapsed. A negative probe was called an outlier if it met one of two criteria. First, if the average count of a probe across all segments was <10% of the average count of all negative probes the probe was removed from all segments. Second, if the probe was called an outlier by the Grubb’s test with alpha = 0.01, it was removed from that segment. If the probe was an outlier by the Grubb’s test in ≥20% of segments, it was removed from all segments. The geometric mean of the remaining probes was calculated to collapse the negative probes to a single count value.

The limit of detection (LOD) above which a gene was called “detected” was defined as 2 standard deviations above the geometric mean of negative probes. For the analyses of the kidney and mouse embryo data, genes were filtered to only those above LOD in >1% of AOIs and counts were normalized by Q3 normalization after removal of genes consistently below LOD. For the CRC/NSCLC differential expression, ssGSEA, and cell type deconvolution analyses, genes were filtered to only those above LOD in >15% of AOIs and counts were normalized by Q3 normalization after removal of genes. For all other datasets and analyses, genes were not filtered and raw deduplicated counts were used.

For the CRC/NSCLC sequencing subsampling analysis, raw FASTQ files were subsampled to the desired read depths using seqtk (https://github.com/lh3/seqtk). Five replicates of the subsampling were performed at each read depth level and all subsamples were run through the NGS data processing pipeline independently. For analyses where sequencing read depth was compared, AOIs were not filtered for sequencing saturation. For analyses where only one read depth is presented, the 150 reads/μm^2^ level was used and AOIs with <50% sequencing saturation were removed from the analysis.

All statistical analyses and data visualizations were performed in R or using the DSPDA software v2.3. Differential expression was performed using a linear mixed effect model with slide and DSP instrument as random effect variables, and p-values were corrected for multiple hypothesis testing. ssGSEA was performed using the GSVA R package (Hänzelmann et al. 2013). Cell type deconvolution was performed using the SpatialDecon R package (Danaher et al. 2022).

## Data Access

All raw and processed data generated in this study are available at the NCBI Gene Expression Omnibus (GEO; https://www.ncbi.nlm.nih.gov/geo/), accession number GSE190089.

## Acknowledgments

We thank Michelle Kriner and Hiromi Sato from NanoString Technologies for reviewing and editing the manuscript.

