## Supplemental Figures for "Spatially resolved whole transcriptome profiling in human and mouse tissue using Digital Spatial Profiling"

A

B

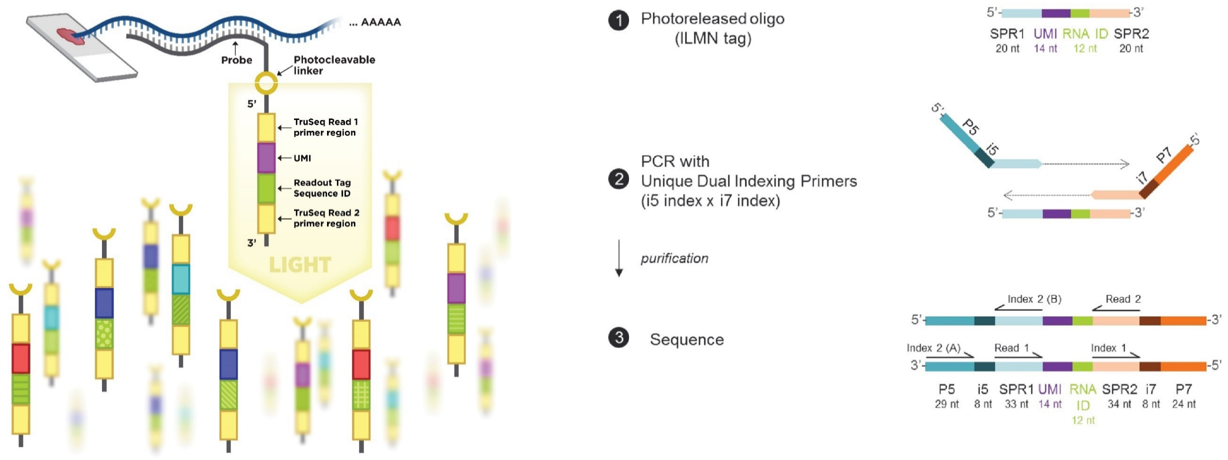


C

| | Human WTA | Mouse WTA |
|---|---|---|
| Number of Genes Targeted | 19505 | 21596 |
| Number of Protein Coding Genes Targeted | 19128 | 21040 |
| Negative Control Probes | 139 | 210 |
| Total Number of Probes | 18815 | 20175 |

**Figure S1. WTA probe design. A.** Schematic of WTA probes, with a sequence complementary to the target RNA, a UV-photocleavable linker, and an indexing tag sequence designed to be read out by next generation sequencing (NGS). The indexing tag sequence contains a UMI and a barcode designed to uniquely identify each probe. **B.** After release on the DSP instrument, the tag sequence is collected and amplified by PCR to add Illumina P5 and P7 sequences and i5 and i7 index sequences for sample demultiplexing after sequencing. **C.** Number of probes and number of genes targeted by the human and mouse Whole Transcriptome Atlas assays.



FOR RESEARCH USE ONLY. Not for use in diagnostic procedures

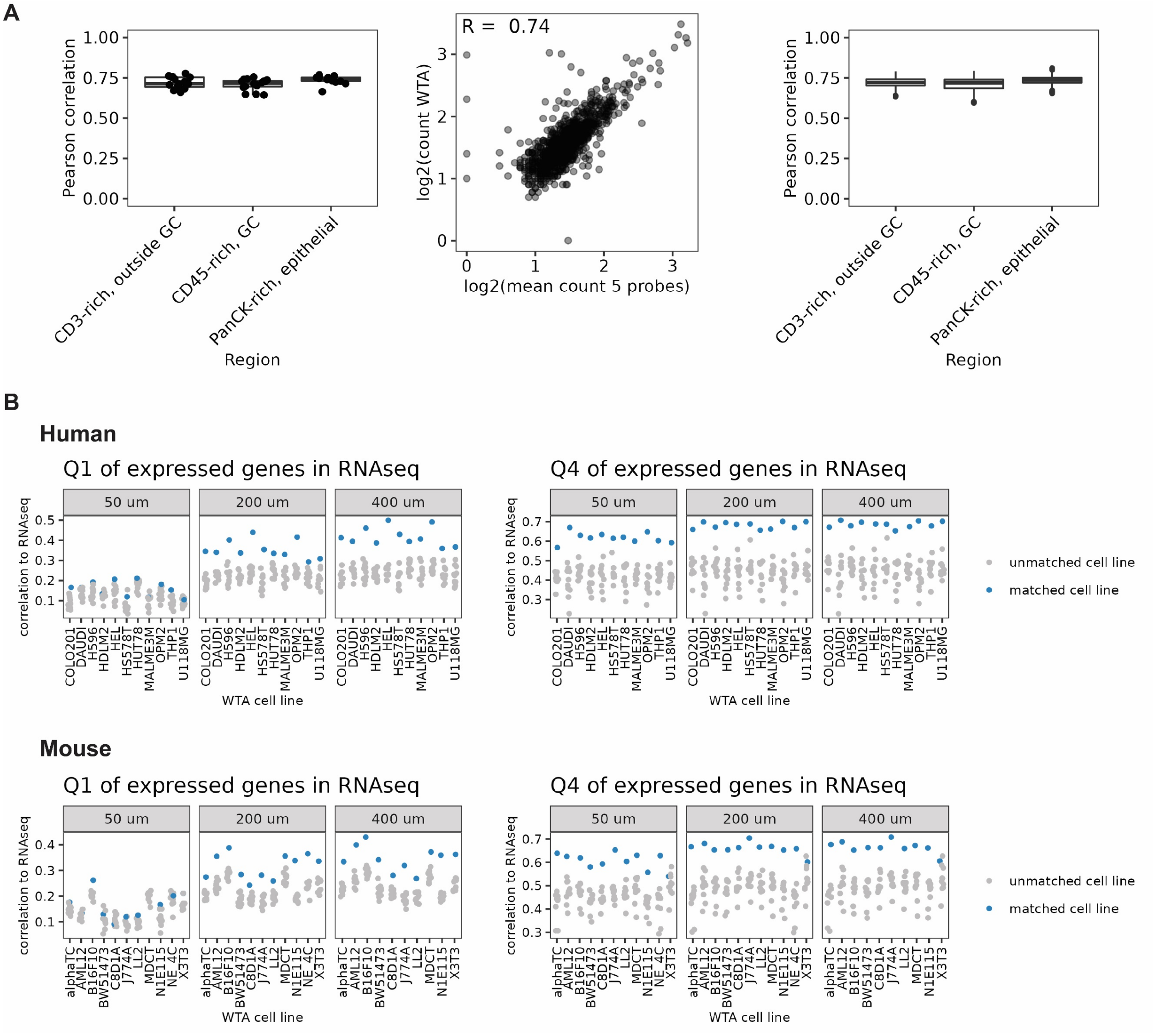


**Figure S2. Correlation of WTA with other DSP panels and RNAseq. A.** Left: scatterplot of WTA counts and the mean counts of an 1812 gene panel with 5 probes per target for a representative matched AOI. Middle/Right: boxplots of Pearson's correlation coefficients between WTA and the mean count of 5 probes per target from the smaller panel, or 100 iterations of a randomly selected single probe. **B.** Spearman's correlation of WTA counts in each AOI compared to RNAseq of each cell line profiled in this experiment using just the lowest quartile of expressed genes in the RNAseq data (Q1), or the highest quartile of expressed genes (Q4). For each AOI, the matching cell line is shown in blue and all other cell lines in grey.

**A**

**Human**

| Gene | H596 | SUDHL4 |
|---|---|---|
| *MS4A1* | 0.07 | 690.15 |
| *PTPRC* | 0.09 | 166.38 |
| *CD27* | 0.19 | 181.92 |
| *CD48* | 0.43 | 218.28 |
| *CD79A* | 0.5 | 656.46 |
| *CXCL8* | 250.77 | 0.59 |
| *DKK1* | 258.89 | 0.4 |
| *C1R* | 350.12 | 0.47 |
| *ITGB4* | 358.74 | 0.93 |

**Mouse**

| Gene | NE-4C | J774A.1 |
|---|---|---|
| *Ptprc* | 0 | 209.415 |
| *Aif1* | 0.106361 | 277.822 |
| *Itgam* | 0.139225 | 451.785 |
| *Rac2* | 0.27408 | 298.922 |
| *Sparc* | 114.693 | 0 |
| *Enpp2* | 116.856 | 0 |
| *Cd63* | 256.868 | 0 |
| *Epcam* | 306.483 | 0 |

**B**

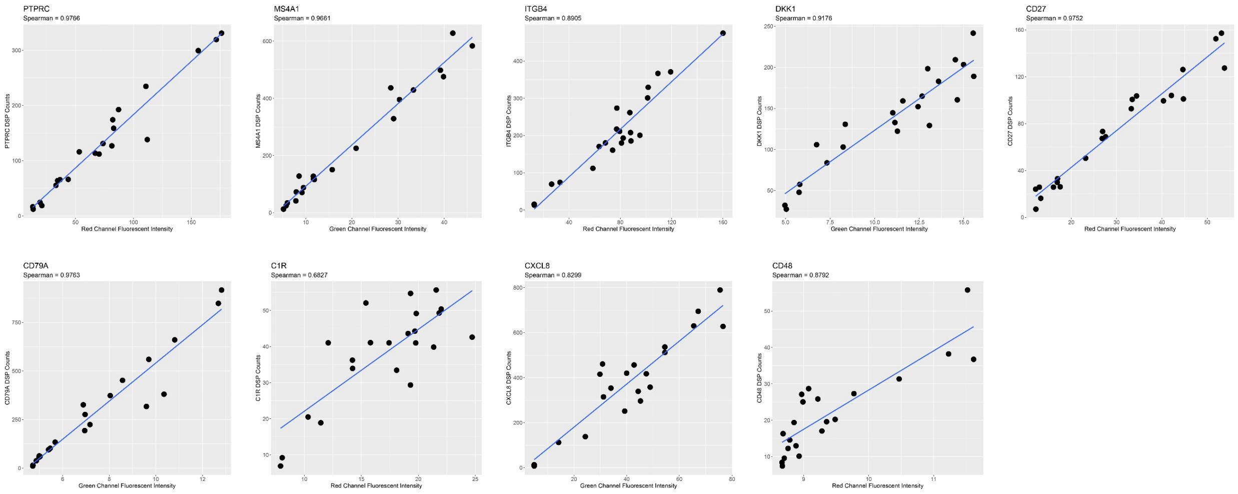


**C**

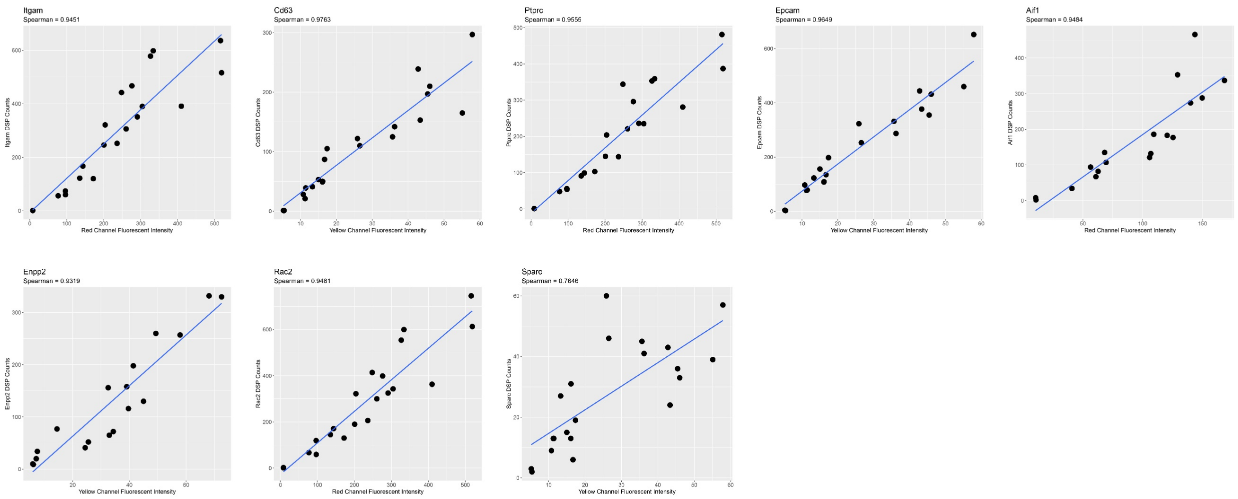


**Figure S3. Correlation of WTA with RNA FISH in cell line titrations. A.** Table showing RNAseq TPM in each of the two cell lines in the titration for each gene tested by RNA FISH. Genes were selected for high (>100 TPM) in one cell line, and low (<1 TPM) expression in the other to create a gradient of gene expression across the titration. **B-C.** Scatterplots showing WTA counts vs RNA

FISH fluorescent intensity across the cell line titration for each gene tested in human (**B**) and mouse (**C**).

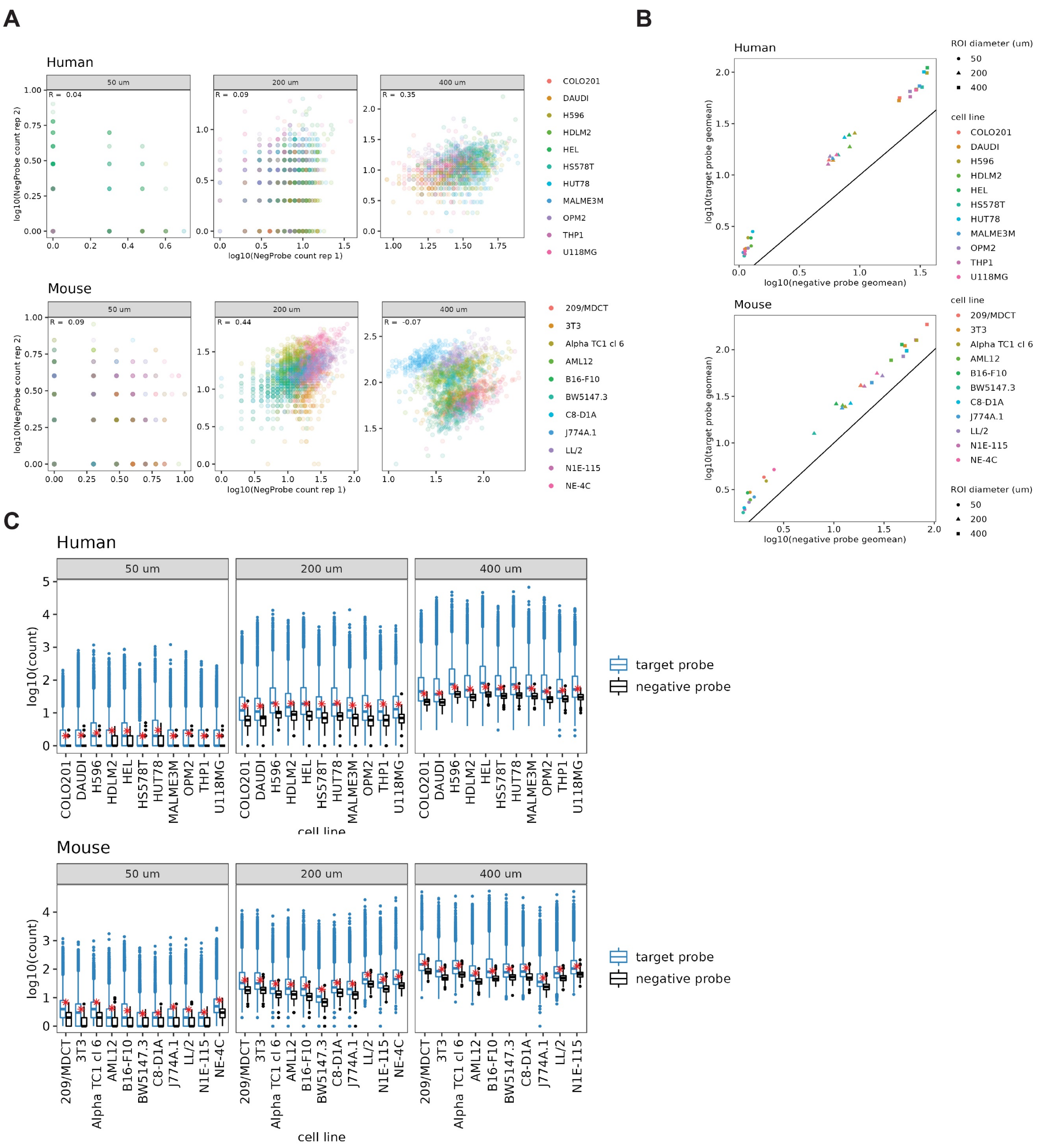


**Figure S4. A.** Log10 counts of all negative probes across replicate experiments in 11 FFPE human and mouse cell lines. Points are colored by cell line, and are separated by AOI size. **B.** Relationship between negative probe signal and target signal in each AOI in 11 FFPE human and mouse cell lines. The log10 geometric mean of negative probes is plotted on the x-axis, and the log10 geometric mean of target probes on the y-axis. Probes were filtered for outliers as described in the Methods. Line is y=x, and points are colored by cell line and shaped by AOI size. **C.** Boxplots showing target (in blue) and negative probe counts (in black) for all genes in 11 FFPE human and mouse cell pellets in 50, 200, and 400 μm diameter circle AOIs. The red dot indicates 2 SD above the geometric mean of negative probe counts, the threshold used for calling a gene above background in the analyses shown in this manuscript.

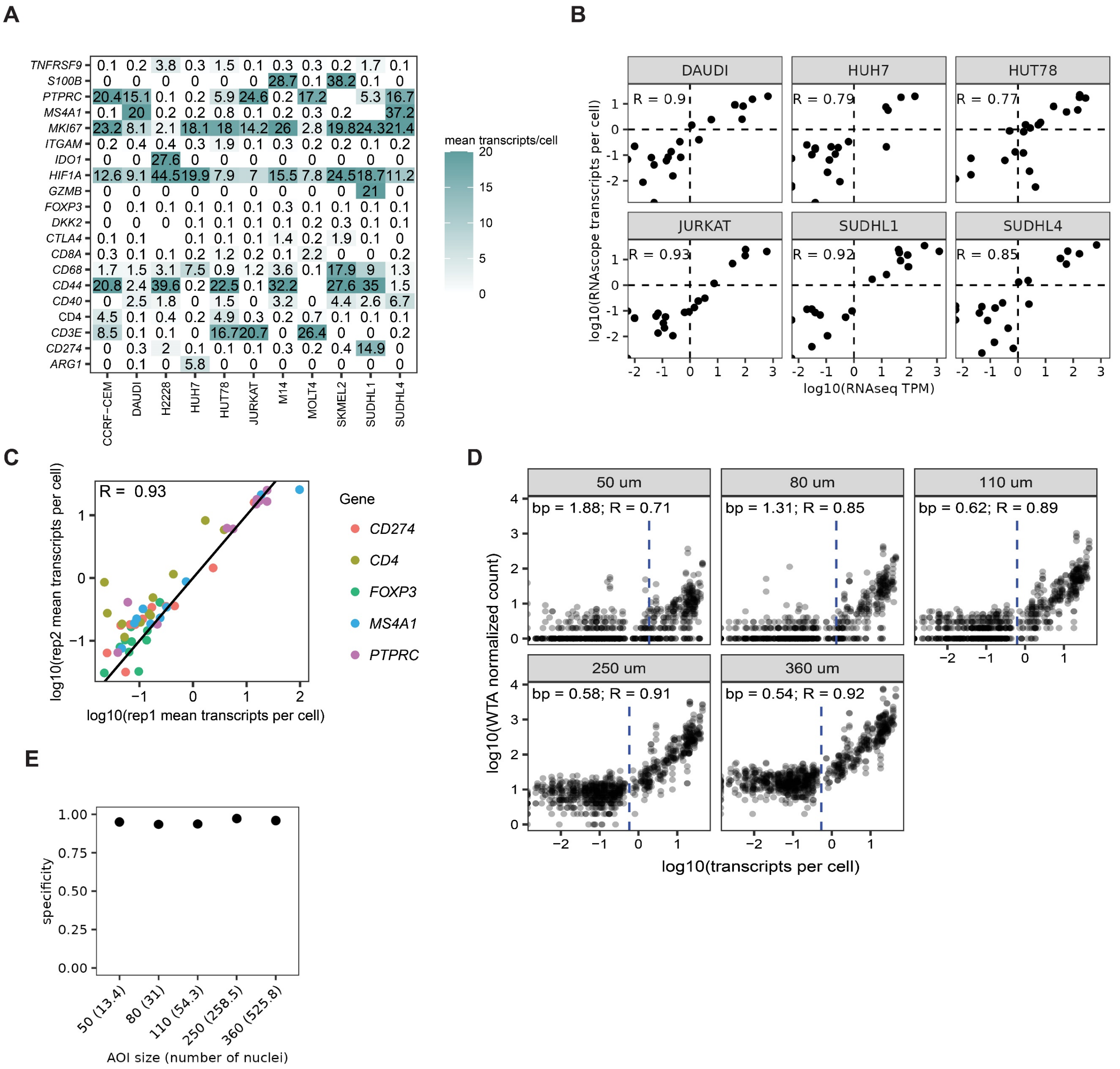


**Figure S5. Sensitivity of WTA at different gene expression levels. A.** Expression level of transcripts used in the RNAscope absolute transcript comparison experiment. Mean number of transcripts per cell for each gene in each cell line is indicated. **B.** Scatterplots plotting log10(TPM) in the RNAseq experiment on the x-axis, and log10(mean transcripts per cell) in the RNAscope experiment for each gene in each cell line tested in this experiment. Dashed lines indicate the threshold for calling a gene "expressed": 1 TPM in RNAseq, and 1 transcript per cell in RNAscope. **C.** Replicability of RNAscope mean transcripts per cell for 5 genes run in duplicate experiments.

**D.** Scatterplot of WTA counts on the y-axis vs mean transcripts per cell from RNAscope on the x-axis. Breakpoints at which WTA counts and RNAscope transcripts per cell become linearly correlated were calculated for each AOI size and are indicated by a dashed line. **E.** Specificity of human WTA compared to RNAscope in each AOI size, using RNAscope mean transcripts per cell ≥ 1 as the threshold for a true expressed gene.

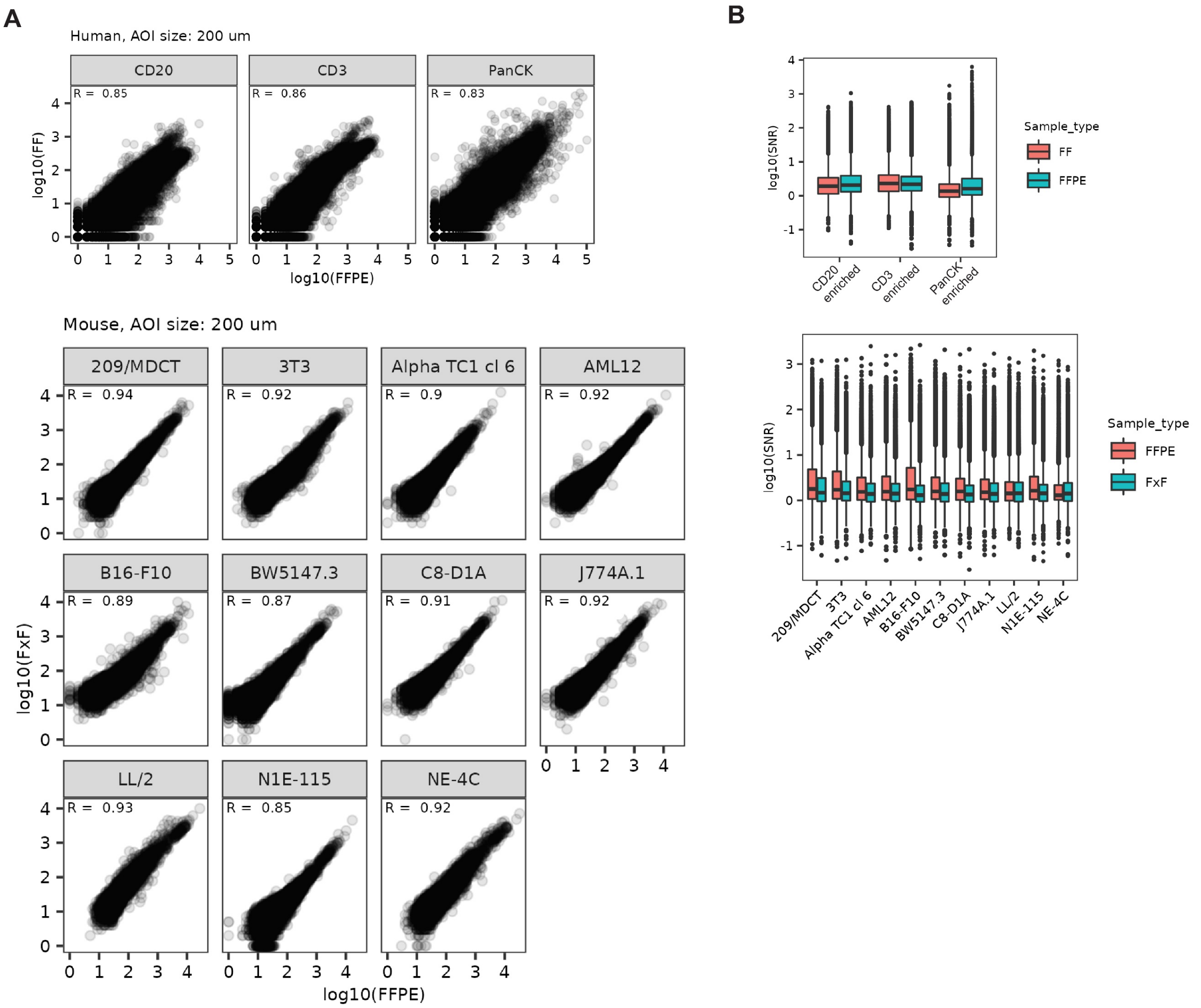


**Figure S6. WTA is compatible with human fresh frozen and mouse fixed frozen samples.** **A.** Scatterplots of WTA counts for matched regions in FFPE vs fresh frozen (FF) tonsil tissue (human, top) and FFPE vs fixed frozen (FxF) cell pellets (mouse, bottom). Pearson correlation coefficients are indicated on each plot. **B.** Comparison of signal-to-background ratio of all genes in FFPE vs FF tonsil tissue (human, top) and FFPE vs FxF cell pellets (mouse, bottom).

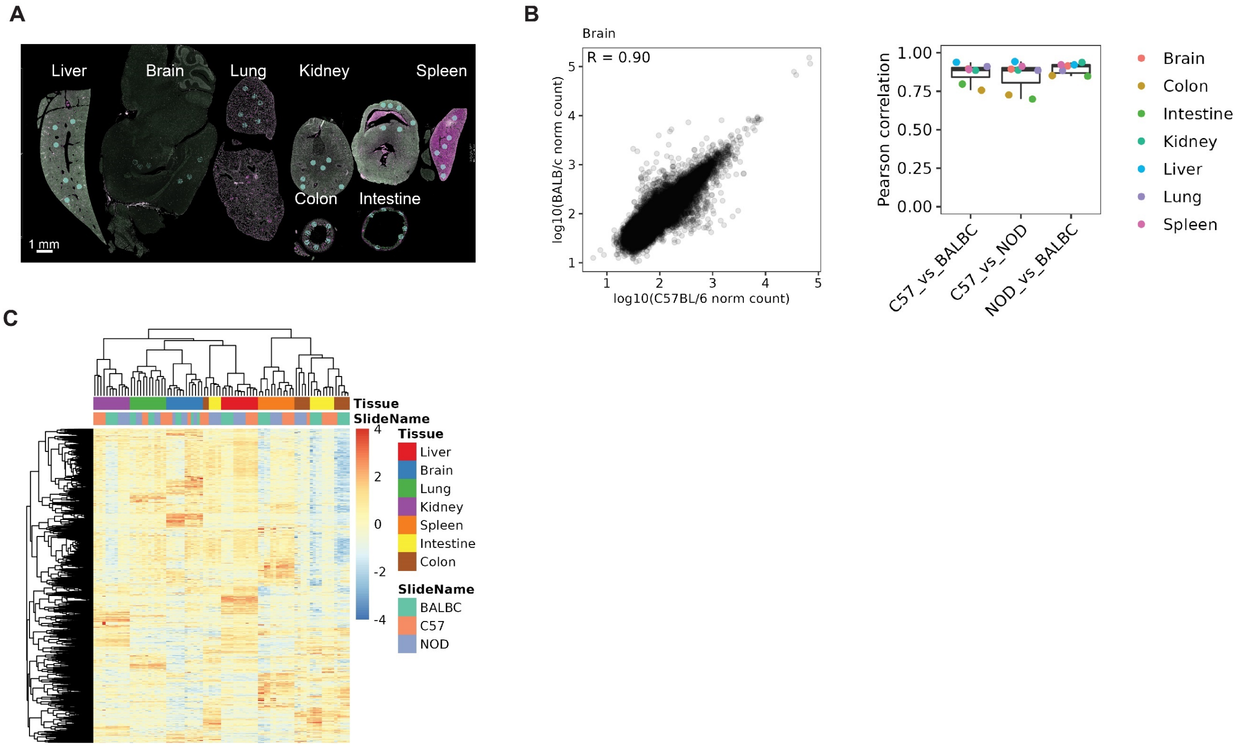


**Figure S7. Mouse WTA is compatible with commonly used mouse strains. A.** Image of the C57BL/6 FFPE organ array used in this experiment, with the 7 tested organs labeled. Similar tissue arrays from two other mouse strains (BALB/c, NOD) were also profiled. **B.** Left: Representative scatterplot of WTA counts in brain between C57BL/6 and BALB/c. Right: Boxplot of correlation coefficients of all comparisons between strains for each organ. **C.** Heatmap of scaled gene expression of the 16610 genes expressed above background in at least 10% of AOIs. AOIs and genes are clustered by hierarchical clustering.

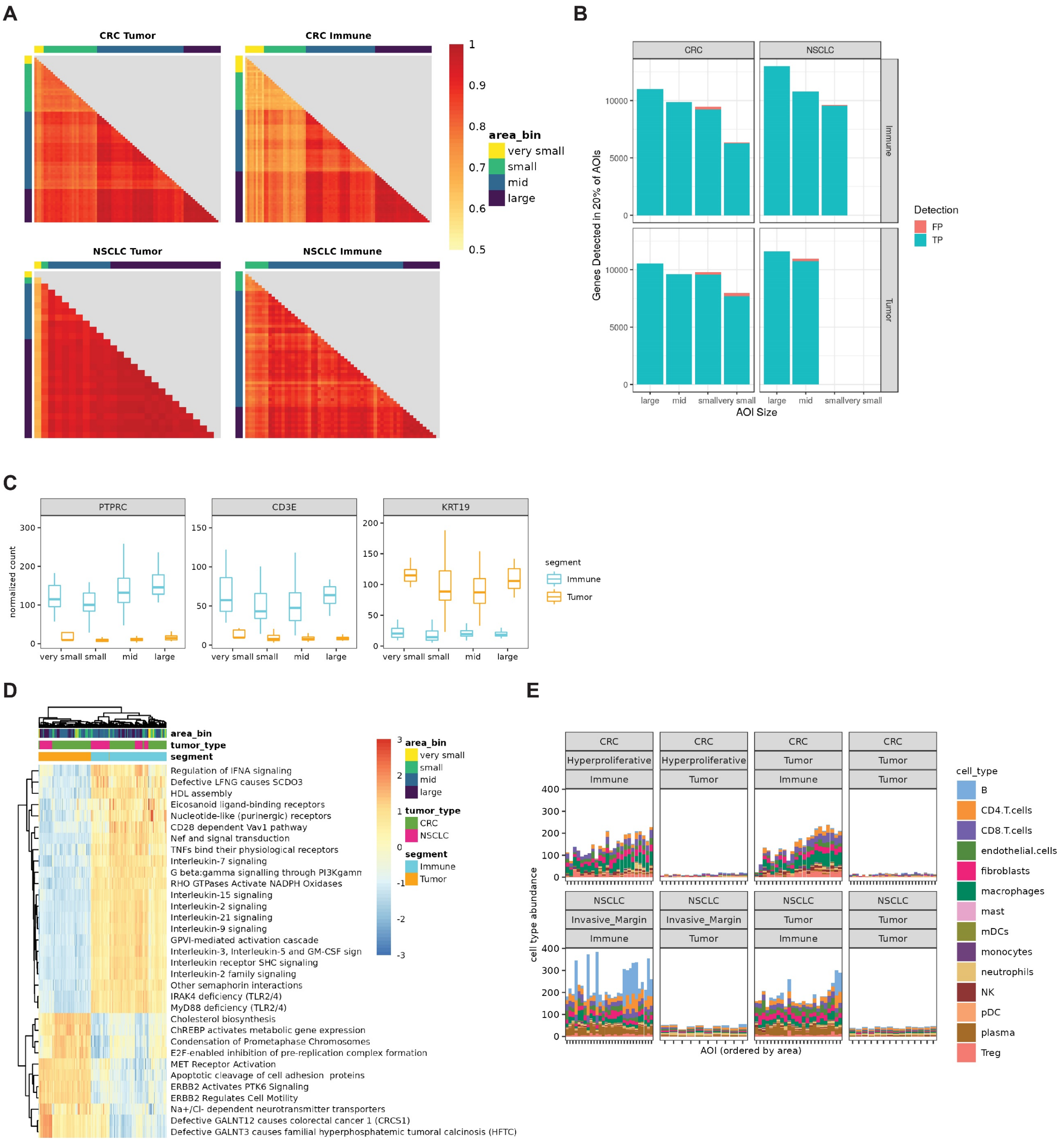


**Figure S8. Read count correlation, genes detected, and cell type deconvolution results across AOI sizes in CRC and NSCLC. A.** Pearson correlation of counts from all genes from each AOI to every other AOI of the same tumor and segmentation type. AOIs are ordered by area. **B.** Number of genes detected in at least 20% of AOIs in each size bin that are also detected in the “large” size bin. Shared genes are in blue, unique genes are in red. **C.** Boxplots showing Q3 normalized count of genes encoding proteins targeted by the antibodies used as morphology markers for the segmentation experiment (CD3E, CD45, and PanCK) in immune and tumor segments binned by area. **D.** Heatmap of ssGSEA enrichment of the most differentially expressed Reactome pathways between tumor and immune compartments. Enrichment was computed using genes detected above background in >20% of AOIs. Columns and rows are clustered by hierarchical clustering and annotated by tumor type, segment, and area bin. All displayed pathways are significant at FDR < 0.01. **E.** Results of cell type deconvolution using a cell profile matrix derived from gene expression profiles of tumor, stroma, and immune cells (Danaher et al.). Data is shown as stacked barplots with each bar as a single AOI and the estimated abundance of each immune cell type colored, and faceted by tumor type, segment, and region within the tumor.

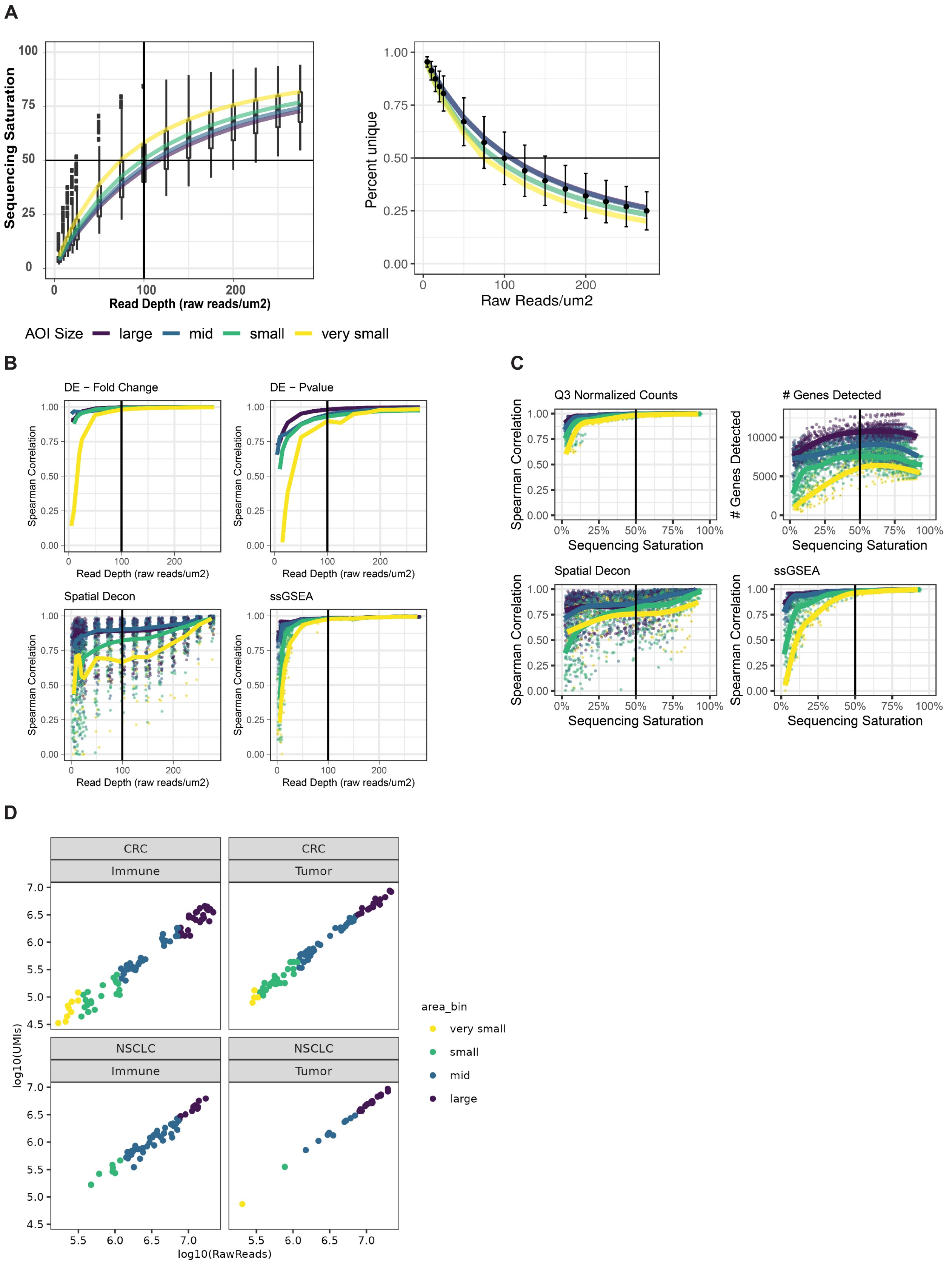
A
Sequencing Saturation
Read Depth (raw reads/um2)
Percent unique
Raw Reads/um2
AOI Size
large
mid
small
very small
B
DE – Fold Change
DE – Pvalue
Spatial Decon
ssGSEA
Spearman Correlation
Read Depth (raw reads/um2)
C
Q3 Normalized Counts
### Genes Detected
Spatial Decon
ssGSEA
Spearman Correlation
### Genes Detected
Sequencing Saturation
D
CRC
Immune
Tumor
NSCLC
log10(UMIs)
log10(RawReads)
area_bin
very small
small
mid
large

**Figure S9. Effect of read depth on performance metrics and biological conclusions from human WTA. A.** Relationship between read depth (raw reads/μm$^2$) and sequencing saturation (1-deduplicated/aligned) (left) or percent unique reads (right) split by AOI size. **B.** The first panel shows genes detected above background per AOI at each size and subsampling level. The next panels show Spearman's correlation of counts, results from cell type deconvolution, differential expression fold change and p-value, and ssGSEA enrichment in subset AOIs to the same AOI at 300 raw reads/μm$^2$. Jittered points are individual AOIs and lines represent the average for each AOI size and are colored by area bin. DE does not have individual AOI points since it is run by treatment (Tumor vs Immune) rather than by AOI. AOI sizes are colored as in A. **C.** Same calculations as B but plotted against individual AOI's sequencing saturation instead of read depth. **D.** Scatterplots showing log10 counts of raw reads versus log10 UMI deduplicated counts for each AOI in this experiment. Data is shown for the 150 reads/μm$^2$ subsampling level.

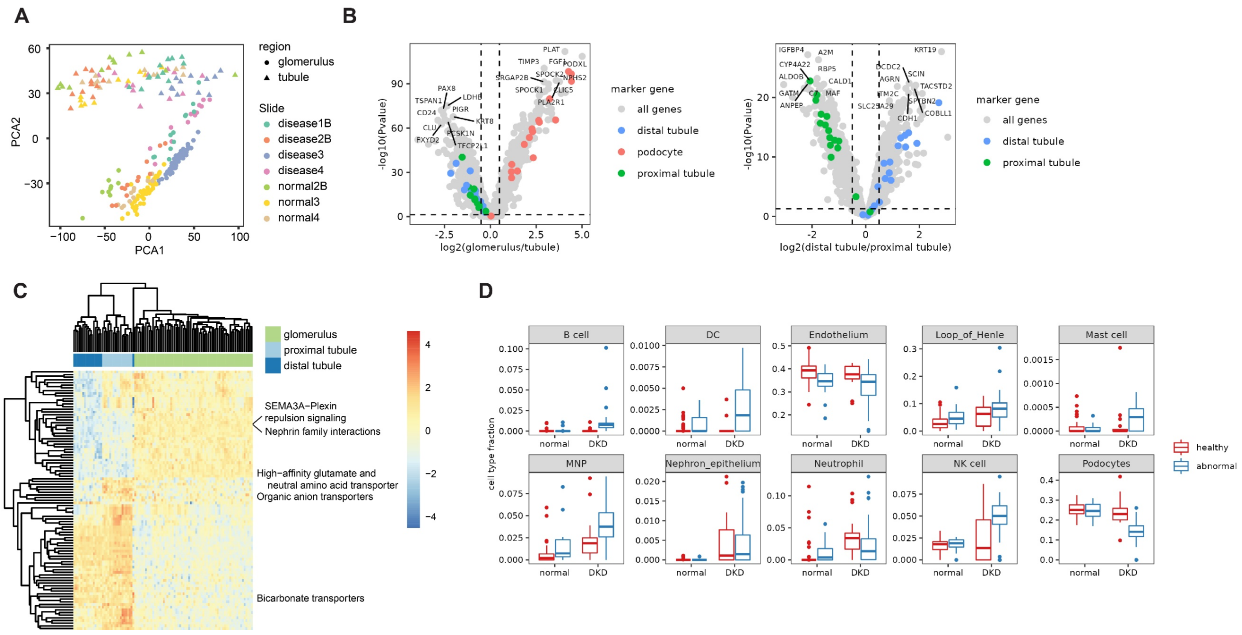


**Figure S10. Additional analyses of the human kidney dataset. A.** Principal component analysis of variation between samples using genes detected above background in >1% of AOIs. PCA1 vs PCA2 is plotted, with points colored by sample and shaped by kidney structure. **B.**

Volcano plots of fold change vs -log10(p-value) from differential expression analysis of normal glomeruli vs tubules, and proximal tubules vs distal tubules. Top marker genes identified in proximal tubule cells, distal tubule cells, and podocytes from single cell RNAseq expression data from (Young et al. 2018) are colored. **C.** Heatmap of ssGSEA enrichment of the most differentially expressed Reactome pathways between substructures in normal kidney samples. Enrichment was computed using genes detected above background in >1% of AOIs. Columns and rows are clustered by hierarchical clustering and the data are scaled by row. All displayed pathways are significant at false discovery rate (FDR) <0.05. **D.** Boxplots of all cell types with significantly different proportions between normal and DKD glomeruli (t-test Bonferroni-corrected p-value < 0.05), colored by whether the glomerulus was annotated as pathologically abnormal or healthy.

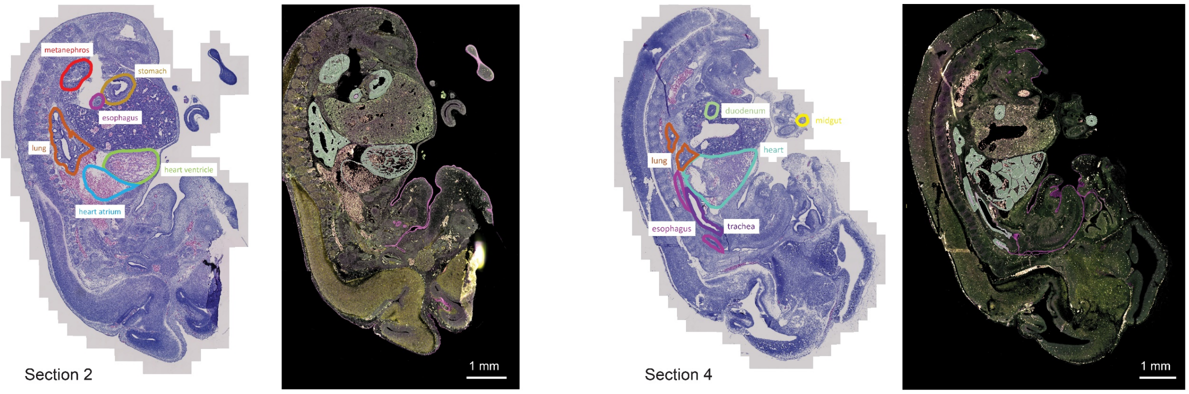


**Figure S11. Annotation of mouse embryo organs.** DSP fluorescent images (left) and H&E images of serial sections (right) of two representative sections of mouse embryo. Organs profiled are outlined and labeled.

#### Supplemental Table Legends

**Supplemental Table 1.** Probe information for all human and mouse WTA probes.

**Supplemental Table 2.** Antibodies and RNAscope probes used in this study.
